# SOX9-regulated matrix proteins predict poor outcomes in patients with COVID-19 and pulmonary fibrosis

**DOI:** 10.1101/2024.01.21.576509

**Authors:** Laurence Pearmain, Elliot Jokl, Kara Simpson, Lindsay Birchall, Yaqing Ou, Craig Lawless, Angela Simpson, Lizzie Mann, Nick Scott, Rajesh Shah, Rajamiyer Venkateswaran, Stefan Stanel, Conal Hayton, Pilar Rivera-Ortega, Phil Hansbro, Neil A Hanley, John F Blaikley, Karen Piper Hanley

## Abstract

Pulmonary fibrosis is an increasing and major cause of death worldwide. Understanding the cellular and molecular mechanisms underlying the pathophysiology of lung fibrosis may lead to urgently needed diagnostic and prognostic strategies for the disease. SOX9 is a core transcription factor that has been associated with fibrotic disease, however its role and regulation in acute lung injury and/or fibrosis have not been fully defined. In this study we apply a hypothesis based approach to uncover unique SOX9-protein signatures associated with both acute lung injury and fibrotic progression. Using *in vivo* models of lung injury in the presence or absence of SOX9, our study shows SOX9 is essential to the damage associated response of alveolar epithelial cells from an early time-point in lung injury. In parallel, as disease progresses, SOX9 is responsible for regulating tissue damaging ECM production from pro-fibrotic fibroblasts. In determining the *in vivo* role of SOX9 we identified secreted ECM components downstream of SOX9 as markers of acute lung injury and fibrosis. To underscore the translational potential of our SOX9-regulated markers, we analysed serum samples from acute COVID19, post COVID19 and idiopathic pulmonary fibrosis (IPF) patient cohorts. Our hypothesis driven SOX9-panels showed significant capability in all cohorts at identifying patients who had poor disease outcomes. This study shows that SOX9 is functionally critical to disease in acute lung injury and pulmonary fibrosis and its regulated pathways have diagnostic, prognostic and therapeutic potential in both COVID19 and IPF disease.

## Introduction

Pulmonary fibrosis is an incurable disease with an average survival of 3-5 years (*1–3*). It is characterised by excessive deposition of extra-cellular matrix (ECM) proteins from activated fibroblasts resulting in scarring and impaired lung function (*4*). Idiopathic pulmonary fibrosis (IPF) is the most common form of the disease and often diagnosed at an advanced stage with variable progression (*5–7*). Identifying patients at an earlier stage and those who are at risk of progression would provide a valuable clinical tool for management of the disease.

Despite progress in diagnostic markers of pulmonary fibrosis, reliable tests capable of predicting progression remain limited. The majority of prognostic markers using patient demographics, physiological lung function and imaging data have only been successful in advanced disease (*8*). As a result, there has been a major interest in understanding the cellular and molecular mechanisms underlying the pathophysiology of lung fibrosis to identify early markers of progressive disease. One common feature associated with the majority of fibrotic lung diseases is increased deposition of ECM proteins (*4, 9*). Consequently identifying core regulators of ECM deposition may be a critical link for identifying downstream targets as biomarkers of progressive disease (*10*).

The transcription factor, Sex determining region Y box 9 (SOX9) regulates many developmental processes, including the proliferation and production of ECM proteins from chondrocytes (*10*). Similarly, we have previously identified SOX9 is a key factor regulating multiple components of the fibrotic ECM (*11–16*). In human, mutations in *SOX9* cause campomelic dysplasia (CD) characterised by a failure of chondrogenesis and infants born with CD often die due to respiratory distress (*10*). In lung development, SOX9 is expressed in undifferentiated epithelial cells to ensure correct branching morphogenesis and is down-regulated in terminally differentiated type 1 and type 2 alveolar cells (AT1 and AT2 respectively) (*17–20*). SOX9 is expressed in a Keratin 5 (KRT5) /SOX9 positive basal airway progenitor population with regenerative capacity (*21–23*). However, in IPF, these healthy cellular regenerative processes are impaired.

AT2 cells capable of self-renewal and AT1 differentiation are lost in IPF resulting in functional decline of the entire gas exchange surface of the lung (*24*). Alongside increased activation of fibroblasts and ECM deposition, single cell sequencing studies of IPF patient samples identified SOX9 in a Keratin 17 (KRT17) disease associated epithelial cell population termed aberrant basaloid cells (*25, 26*). Significantly, these disease associated SOX9+/KRT17+ cells lack KRT5, potentially as part of the impaired regenerative response. Understanding the role and regulation of SOX9 in early transitioning cellular events following injury and fibrosis is critical to identify downstream targets as early markers of progressive disease.

In this study we utilise experimental models of lung injury following SOX9-loss to uncover the unique protein signatures associated with both acute lung injury and fibrotic progression. To underscore the translational potential of our SOX9-markers, we utilise IPF human tissue and serum samples from acute COVID-19, post COVID-19 and IPF patient cohorts. Our hypothesis driven SOX9-panels showed significant capability in all cohorts at identifying patients who had poor disease outcomes.

## Results

### SOX9 expression is increased following acute lung injury and fibrosis

To understand the role and regulation of SOX9 in lung injury and fibrosis, we first determined the spatiotemporal expression of SOX9 during acute lung injury through to fibrosis from bleomycin-induced damage (Fig.1A) (*27*). SOX9 expression was evident as early as 24 hours following bleomycin-induced lung injury with more widespread expression and evolving spatial patterns, throughout the course of progressive damage and fibrosis (Fig.1B). During acute injury (days 1-3), SOX9 was mainly located proximal to injured airways and vasculature, alongside occasional alveoli (Fig.1B). Whereas, mice given saline control only showed SOX9 expression in basal cells of proximal airways, in keeping with known expression in health (Fig.1B) (*28*). At days 7-14 (inflammatory-fibrosis time points) increasing stroma and developing fibrotic architecture was observed (dashed line, Fig.1B), with SOX9 positive cells present on the edge of organising parenchyma (Fig.1B) and proximal to alveoli (Fig.1B, C). Dual immunohistochemistry (IHC) confirmed these findings, showing SOX9 localisation with CD31+ vasculature and PDGFRβ+ stroma throughout the injury time-course (Fig.1C and Fig.S1). Picosirius red staining (PSR) identified established collagen deposition at day 28, (Fig.1D). Sequential sections identified SOX9 expression localised to regions of extracellular matrix (ECM) deposition (PSR), as well as the fibroblast marker α-SMA (Fig.1D).

**Fig. 1.**
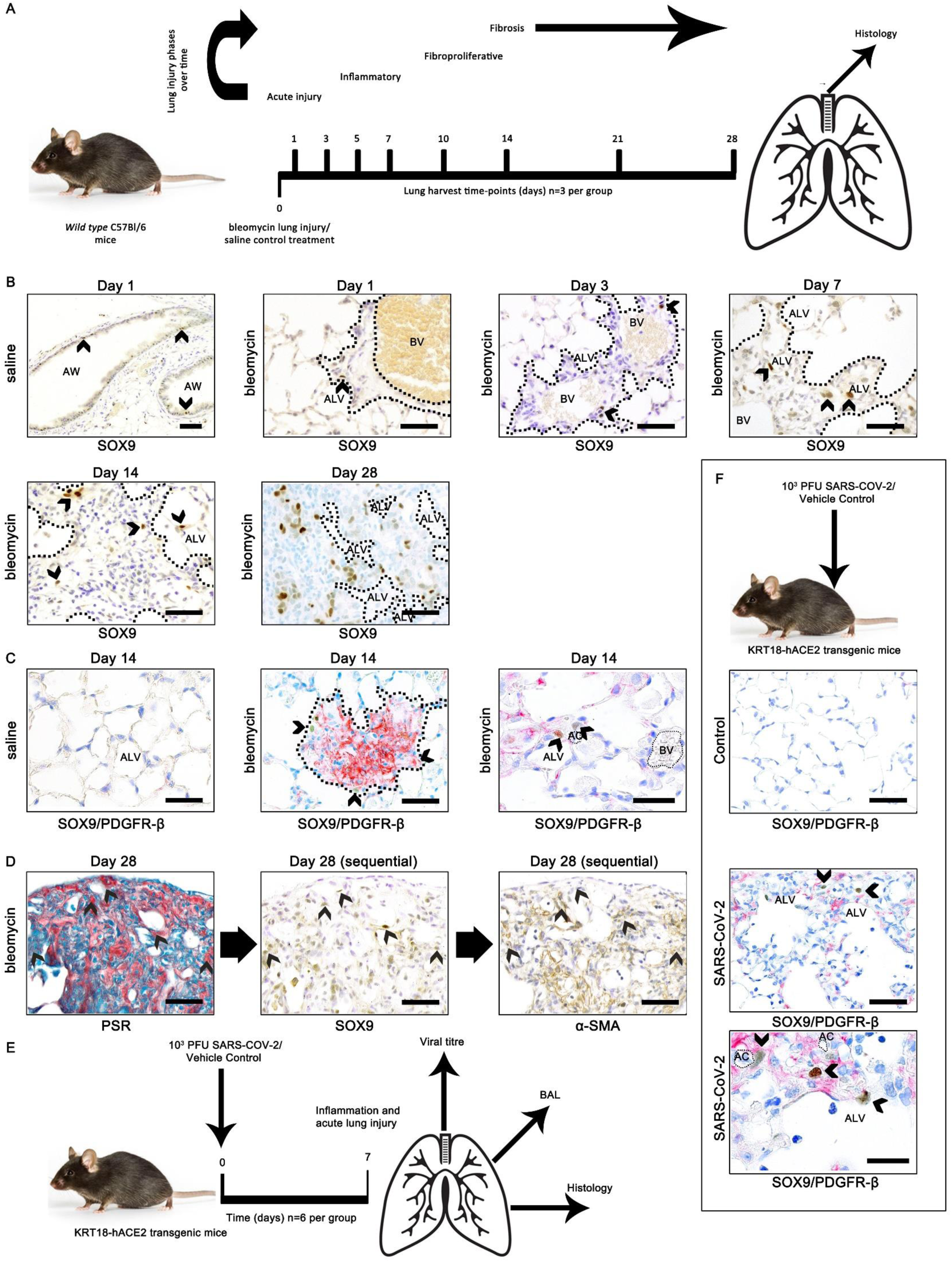
*Wild type* bleomycin time-course and KRT-18hACE2 transgenic SARS-Cov-2 models showing SOX9 temporal and spatial expression. (**A**) Diagram of bleomycin lung injury time-course experiment (n=3/group/time-point). (**B**) Immunohistochemistry (IHC) showing SOX9 expression (DAB, brown; black arrows) in saline control (top left) and bleomycin treated mice across time-points. Dashed lines (black) highlight stroma disrupting normal lung architecture. (**C**) SOX9 expression (DAB, brown) and PDGFR-β expression (red) co-localisation of SOX9 in bleomycin treated lung (middle, right) and saline treated (left). High magnification IHC (right) of SOX9 and PDGFRβ co-localisation in bleomycin injured alveolar compartment. AC (alveolar capillaries); ALV (alveoli). (**D**) Sequential sections of bleomycin treated (28 day) lungs. Left image shows collagen by picosirius red (PSR) staining. Middle image shows SOX9 expression (brown, DAB). Right image shows α-SMA expression (DAB, brown). Black arrows denote the same region across sections. Scale bars, 50μm; except high magnification, 20μm. Counterstaining toluidine blue. (**E**) Diagram of KRT-18-h-ACE2 mouse SARS-CoV-2 infection model experiment (n=6/group). (**F**) SOX9 (DAB; brown) and PDGFR-β (red) expression in control (top) and SARS-CoV-2 infected mouse lung (middle). High magnification image shows SOX9 and PDGFR-β co-expression in SARS-CoV-2 injured alveoli (bottom). Abbreviations: AW, airway; BV, blood vessel; ALV, alveolus; AC, alveolar capillary.

As a second model, and to confirm our findings following bleomycin-induced lung injury, we investigated SOX9 expression in a SARS-CoV-2 mouse model (Fig.1E). KRT18-hACE2 transgenic mice infected with 10^3^ PFU SARS-CoV-2 have been shown to replicate features of human severe acute COVID-19 at 7days post infection (*29–32*). SOX9 expression was evident in SARS-Cov-2 infected lungs compared to control and co-localised with PDGFRβ stromal cells in areas of alveolar damage and architectural destruction (Fig.1F).

### SOX9 is important to epithelial dysregulation, its loss protects against acute lung injury

To investigate the functional significance of SOX9 expression during the acute phase of lung injury we utilised a tamoxifen-(Tam) inducible model of global SOX9 loss (Sox9^fl/fl^;ROSACreER^+/−^; *Sox9 KO)* to eliminate SOX9 prior to injury (Fig.2A), contrasting to controls (Sox9^fl/fl^;ROSACreER^-/-^; *WT*) (*33, 34*).

**Fig. 2.**
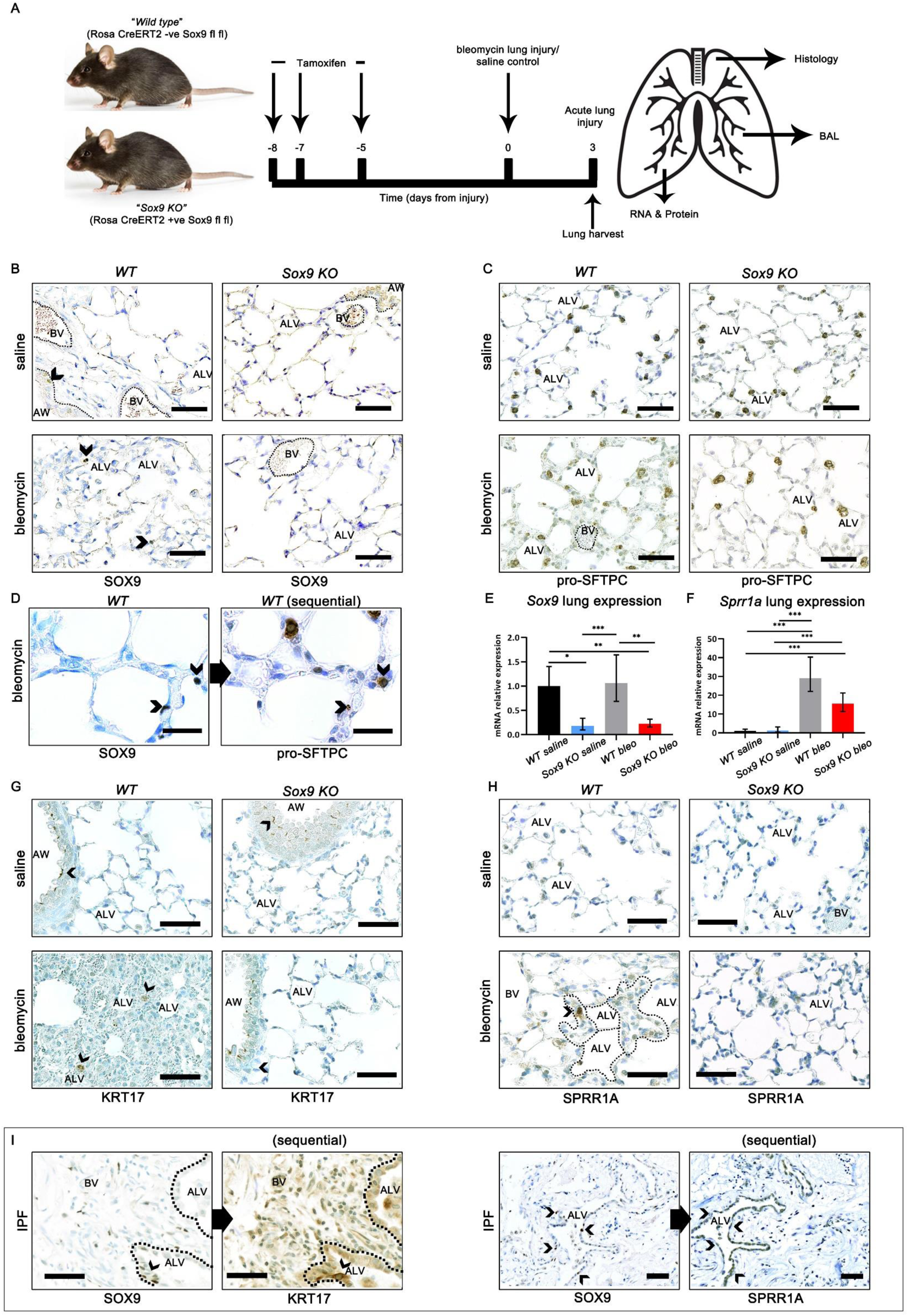
*Sox9 KO* regulates expression of damage associated epithelial populations in acute lung injury, and SOX9 is co-expressed in these populations in fibrotic human lung. (**A**) Diagram of *Sox9 KO* acute lung injury bleomycin model experiment design. (**B**) Immunohistochemistry (IHC, N=3/group) showing SOX9 expression (DAB, brown) of *Wild type* (*WT*; left) and *Sox9 Knockout* (*Sox9KO*; right) lungs treated with saline (top) and bleomycin (bottom). (**C**) Pro-Surfactant Protein C (pro-SFTPC) expression (DAB, brown) shows an increase of pro-STPC+ alveolar cells in *WT* (bottom left), but not *Sox9 KO* (bottom right) bleomycin compared to saline treatment (top row) animals. (**D**) SOX9 (DAB, brown; left) and pro-SFTPC expression (DAB, brown; right) in high magnification images of sequential sections in alveoli of *WT* bleomycin treated lung. (**E**) *Sox9* and (**F**) *Sprr1a* gene expression (mRNA) in whole lung. Significance. (p<0.05) tested using two way ANOVA. (**G**) Keratin 17 (KRT17) expression (DAB, brown) shows an increase of KRT17+ cells in distal airspaces of *WT* (bottom left), but not *Sox9 KO* (bottom right) bleomycin compared to saline treatment (top row) animals. (**H**) SPRR1A expression (DAB, brown) shows an increase of SPRR1A+ cells in distal airspaces of *WT* (bottom left), but not *Sox9 KO* (bottom right) bleomycin compared to saline treatment (top row) animals. (**I**) IHC of human IPF lung showing SOX9 (DAB, brown; left) and KRT17 (DAB, brown) expression in sequential sections and SOX9 (right; DAB, brown) and SPRR1A expression. Black arrows show the same area of damage associated epithelium. Scale bar, 50μm, apart from high magnification, 20μm. Counterstaining with toluidine blue. Abbreviations: AW, airway; BV, blood vessel; ALV, alveolus.

SOX9 expression was upregulated in *WT* bleomycin treated mice, with increased cellular density (indicated by more toluidine blue stained nuclei), compared with the *Sox9 KO* bleomycin group (Fig.2B). The AT2 marker pro-surfactant protein C (pro-SFTPC) expression was also increased (Fig.2C) and SOX9 co-localised to a proportion of pro-SFTPC expressing cells in *WT* bleomycin injured lungs (Fig.2D). In line with previous studies on *Krt17+* transitional alveolar epithelial cell population with a pro-fibrotic gene signature in human IPF lungs (termed aberrant basaloid cells) (*25*), we identified occasional KRT17+ cells in distal airspaces of *WT* bleomycin treated animals.

These KRT17+ cells were not seen in *Sox9 KO* animals in the acute lung injury model (Fig.2G). Moreover, from sequential sections of *WT* fibrotic lung (at 14 day post-bleomycin treatment) we noticed KRT17, SOX9 and SFTPC expression were co-localised in the same areas of alveolar damage proximal to fibrotic regions of lung (Fig.S2B). To determine whether these cells were damage associated transitional epithelial cells, we showed co-localisation of SOX9 and SPRR1a in sequential sections following bleomycin-induced injury (Fig.S2A) (*35*). These co-localisation data were similarly replicated in our SARS-CoV-2 model (Fig.S2C). Similarly to KRT17, there was an absence of SPRR1A in the distal airspace of bleomycin injured *Sox9 KO* animal lungs compared to *WT* in acute injury (Fig.2H). Upregulated expression of SOX9 and co-localisation with KRT17 and SPRR1A was reproducible in alveoli of human IPF lung tissue (Fig.2I). Collectively these data suggest aberrant expression of SOX9 during lung injury has a function in driving the damage associated transitional alveolar epithelial cell response (*25, 35*).

In addition to observed alveolar epithelial changes assessment of bronchoalveolar lavage fluid (BALF) showed *Sox9* KO animals had significantly lower total protein content compared to WT control (Fig.S3A). Immune response following acute injury severity showed *Sox9 KO* animals had fewer leucocytes in the bronchoalveolar space compared with *WT* animals (Fig.S3B, C), indicating a reduced immune response. This was further supported by fewer neutrophils (CD64-/MERTK-/LY6G+; Fig.S3C), NK cells (CD64-/MERTK-/Ly6G-/SiglecF-/CD3-/NK1.1+; Fig.S3D) and monocyte derived macrophages (CD64+/MERTK+/Ly6Chi; Fig.S3E) due to *Sox9 KO*. Collectively, these data suggest *Sox9* loss protects against acute lung injury severity.

### SOX9 loss protects against pulmonary fibrosis *in vivo*

In addition to acute injury, our data also highlighted SOX9 during the fibroproliferative and fibrotic time-points (Fig.1B). To provide mechanistic insight into the role of SOX9 specifically during these phases, and thereby differentiate from effects induced during acute injury, we first induced fibrosis using bleomycin (or saline control) in *Sox9 KO* and WT mice, then administered Tam to delete *Sox9* on days 5, 6 and 8 days following injury and harvested lungs at day 14 (Fig.3A). This lung injury window is most recommended for translational targeting (*27*).

**Fig. 3.**
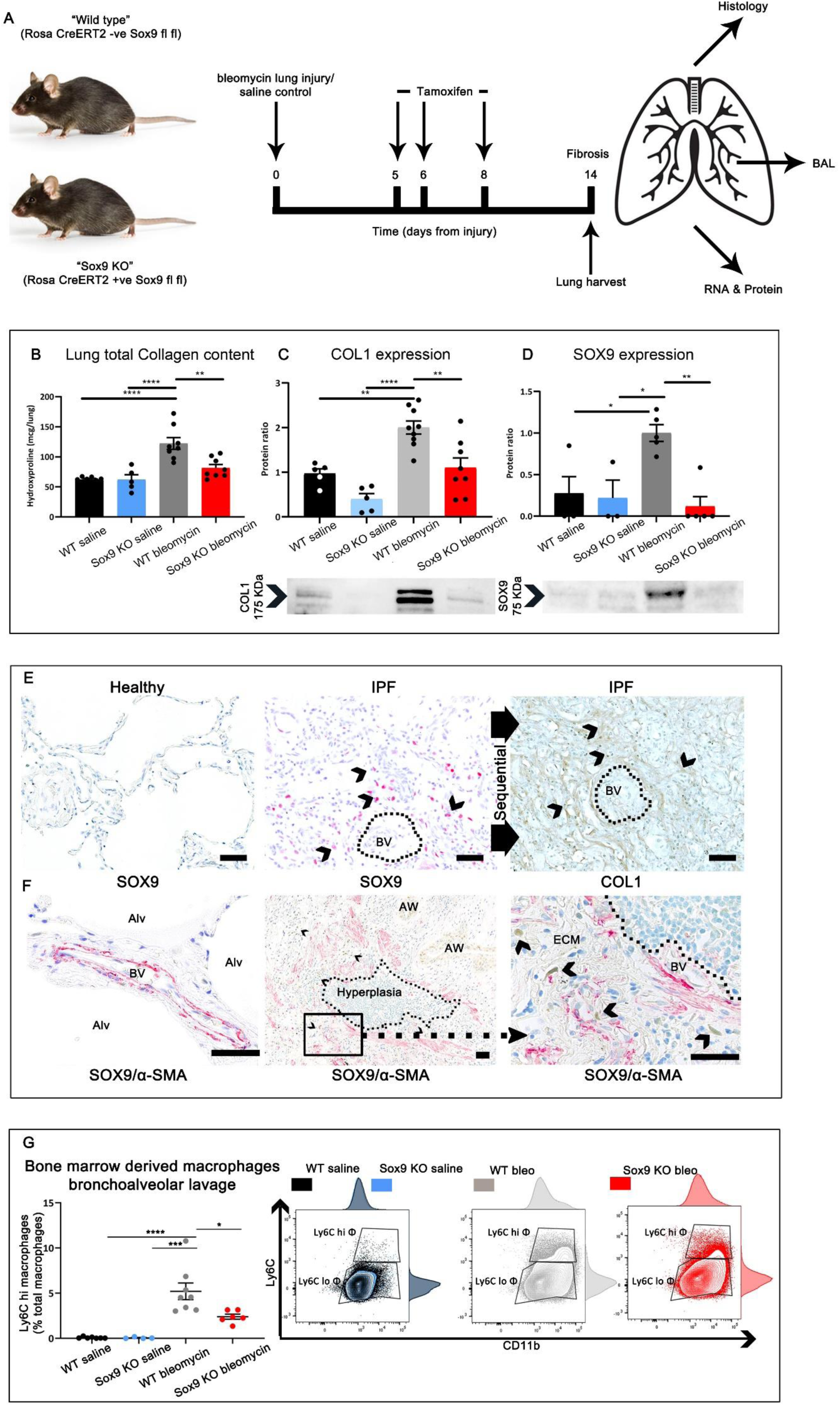
SOX9 regulates extracellular matrix deposition in mouse and human pulmonary fibrosis. (**A**) Diagram of *Sox9 KO* fibrosis bleomycin model experiment (14 days treatment). (**B**) Hydroxyproline for total collagen, per lung, of *Wild type (WT) and Sox9 knockout (Sox9 KO)* saline (n=5/group) and bleomycin (n=8/group) treated animals. (**C**) COL1 protein expression in whole lung from *Sox9 KO* fibrosis experiment animals. (**D**) SOX9 protein expression in *Sox9 KO* fibrosis experiment animals. Statistical significance (p<0.05) of *in vivo* experiments tested by two-way ANOVA (**E**) SOX9 expression (red) in healthy lung (left) and IPF lung (middle), sequential section (right) shows COL1 expression. (**F**) SOX9 (DAB, brown; black arrows) and α-SMA (red) expression in healthy lung (left) and at the edge of fibrosis in IPF (middle, higher magnification of black box region on right). Counterstaining toluidine blue. Scale bar, 50μm. (**G**) Proportion of macrophages (TER119-CD45+CD64+MERTK+) derived from monocytes (TER119-CD45+CD64+MERTK+Ly6Chi), in bronchoalveolar lavage fluid (BALF) from *Sox9 KO* fibrosis experiment animals. Abbreviations: AW, airway; BV, blood vessel; ALV, alveolus; ECM, extra-cellular matrix.

In keeping with SOX9 playing a role in excessive ECM deposition during fibrosis, loss of *Sox9* resulted in reduced pathological collagen assessed by whole lung hydroxyproline assay and COL1 western blot (Fig.3B-D). Overall these findings are consistent with *Sox9* loss reducing the extent of pulmonary fibrosis during the fibroproliferative phase. Moreover, our mouse lung injury time-course model showed SOX9 was localised to regions of ECM deposition and the fibroblast markers PDGFRβ and α-SMA (Fig.1C). This co-localisation was also observed in human control and IPF lung tissue where SOX9 was detected alongside collagen (COL1; Fig.3E) and within myofibroblasts (α-SMA; Fig.3F).

Similar to our acute model (Fig.S3), we assessed immune cell populations in BALF samples (Fig.S4). Total leukocytes (TER119-/CD45+) were reduced in *Sox9 KO* bleomycin animals compared with *WT* (Fig.S4A). Further, monocyte derived macrophages (CD64+/MERTK+/Ly6Chi), as a proportion of alveolar macrophages, were again reduced with *Sox9 KO* in bleomycin treated lungs (Fig.3G). Monocyte derived macrophage accumulation and dysregulation is a crucial component of the fibrotic niche and drives fibrosis (*36, 37*). Further, *Sox9 KO* in the fibrosis model reduced the adaptive immune response, with lower numbers and proportions of lymphocytes (CD4+ and CD8+; Fig.S4C). These data suggest SOX9 indirectly shapes the immune response within the fibrotic alveolar compartment, supported by the lack of SOX9 immune expression across our studies (*33, 34*).

### Identification of biomarkers that identify lung injury progression

*Sox9* ablation both prior to acute injury and during fibroproliferation reduced disease severity and key features of the fibrotic niche. We next aimed to target downstream SOX9 regulated pathways to identify patient relevant biomarkers for lung disease.

Protein was isolated from the supernatant of BALF in animals from both acute lung injury and pulmonary fibrosis *Sox9 KO* and *WT* control mice (Fig.2A and Fig.3A). Unbiased whole proteome analysis was performed. Within the κ-means clusters there was enrichment for terms related to ECM, ECM binding and ECM regulation in both the *Sox9 KO* acute lung injury, as well as fibrosis, models (Fig. 4A-B and E-F). Consistent with the previously presented results proteins within these ECM annotated clusters had lower expression in *Sox9 KO* animals (Fig.4C, G).

**Fig. 4.**
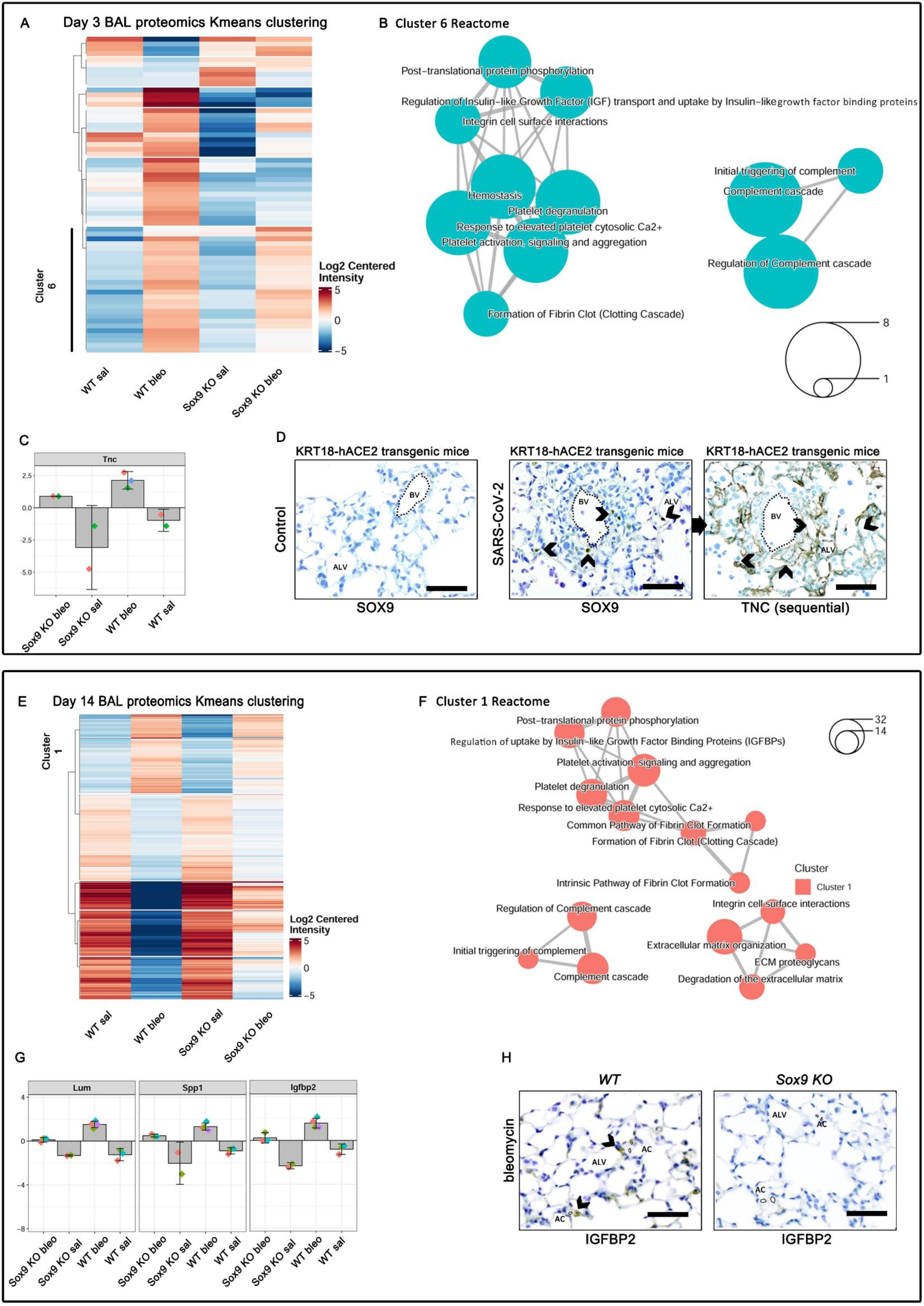
Identification and validation of potential biomarkers downstream of SOX9. (**A-C**) represent results from *Sox9 KO* acute lung injury (3 day treatment) experiments, n=2-3/group. (**A**) Bronchoalveolar lavage (BAL) κ-means cluster whole proteome analysis, detected by liquid chromatography mass-spectrometry. (**B**) cluster 6 reactome map with gene ontology (GO) enriched annotation terms of differentially expressed proteins. (**C**) log_2_ centred intensity of TNC biomarker in *wild type (WT)* and *Sox9 Knockout (Sox9 KO)* animals treated with saline or bleomycin (3 days). (**D**) SOX9 (middle) and TNC (right) expression in sequential sections (DAB, brown) in KRT18-hACE2 control (left) and SARS-Cov-2 lung (middle, right). Black arrows represent the same area in sequential sections. Counterstain toluidine blue. Scale bars, 50μm. (**E-H**) represent results from *Sox9 KO* fibrosis (14 day treatment) experiments. n=2-3/group. (**e**) Bronchoalveolar lavage (BAL) κ-means cluster analysis of whole proteome. (**F**) cluster 1 reactome map with gene ontology (GO) enriched annotation terms. (**G**) log_2_ centred intensity of selected fibrosis progression biomarkers in *Sox9 KO* fibrosis model animals. (**H**) IGFBP2 expression (DAB, brown; black arrows) in *WT* (left) and *Sox9 KO* (right) bleomycin treated. Counterstain toluidine blue. Scale bars, 50μm. Abbreviations: BV, blood vessel; ALV, alveolus; AC, alveolar capillary.

Tenscin C (TNC) and tissue inhibitor of metallopeptidase 1 (TIMP1) were identified from the acute lung injury *Sox9 KO* model and TNC, Lumican (LUM), Osteopontin (SPP1) and Insulin growth factor binding protein-2 (IGFBP-2) from the fibrosis *Sox9 KO* models (Fig.4). As validation, BALF proteomic expression profiles and protein co-localisation with SOX9 were confirmed by IHC in KRT18-hACE2 SARS-CoV-2 and *Sox9 KO* acute lung injury and fibrosis animal models (Fig.4D, H).

As secreted ECM proteins, we hypothesised our SOX9-panel could be useful biomarkers of lung damage and fibrosis progression in patient serum samples. To that end we recruited patient cohorts to assess prediction of severe acute COVID-19 (Cohort 1), diagnosis of “long COVID” related lung damage (Cohort 2), and lung fibrosis progression (Cohort 3).

### Biomarkers identified downstream of SOX9 identifies patients who develop severe acute COVID-19 lung injury

Early identification of severe acute lung injury is critical, as successful therapeutic intervention has a diminishing likelihood of success as injury and acute respiratory distress syndrome progresses (*38*). Identifying a patient’s risk of developing severe acute COVID-19 on admission would act as an early indicator for intensifying subsequent care. To this end we assessed our acute injury markers TNC and TIMP1 in a cohort of patients hospitalised with COVID-19 pneumonia due to SARS-CoV-2 infection. Serum samples were taken at admission to hospital and compared to healthy controls (Cohort 1, Fig.5A). Severity of COVID-19 was classified according to maximal respiratory support requirements as mild, moderate and severe (Fig.5A). Demographics and clinical outcomes can be seen in Table S1.

**Fig. 5.**
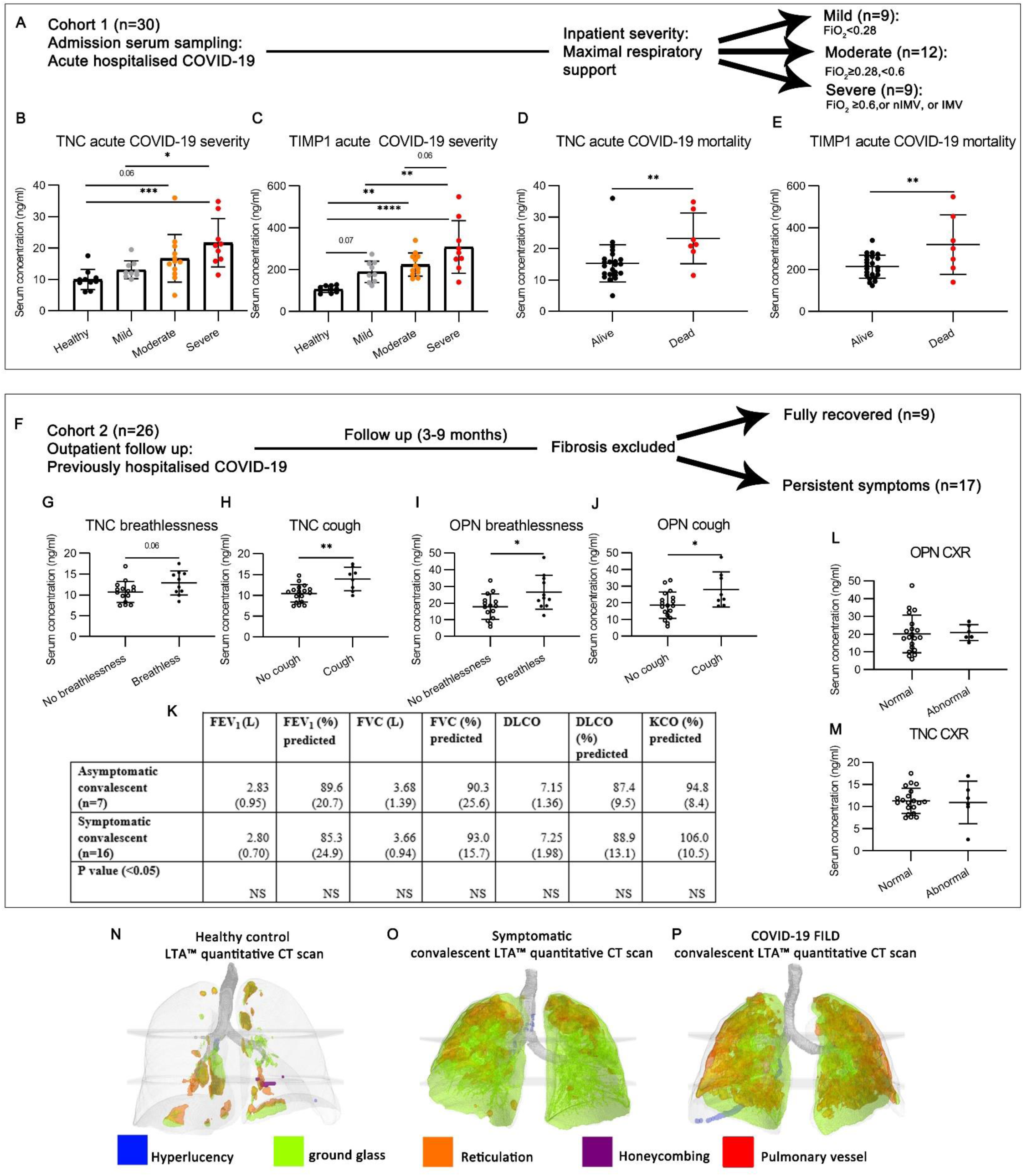
SOX9 biomarkers predict poor outcome in acute hospitalised SARS-CoV-2 infection and identify persistent symptoms in convalescence. (**A**) Cohort 1 (acute hospitalised COVID-19) outcome groups. Serum concentration of biomarkers, at admission, of (**B**) Tenascin C (TNC) and (**C**) TIMP1 (ng/ml) stratified by eventual illness severity. Statistical significance (p<0.05) was tested by one way ANOVA. (**D**) TNC expression and (**E**) TIMP1 serum concentration (ng/ml) at admission, stratified by mortality (7/32). Statistical significance (p<0.05) was tested by two-way unpaired t-test. (**F**) Cohort 2 (convalescent COVID-19, excluding fibrosis) outcome groups. (**G-J**) Serum TNC and Osteopontin (OPN) concentration (ng/ml), at outpatient review, stratified by respiratory symptoms. Statistical significance (p<0.05) tested by two way t-test. (**K**) Table showing lung function performed on non-fibrotic convalescent patients hospitalised with COVID-19, stratified by radiological outcome. Statistical significance (p<0.05) tested by one way ANOVA. Units of measurement: FEV_1_, litres/min; FVC, litres; millilitres CO/minute/mm Hg. % predicted is a comparison to the GLI (2017) reference values. Values in () represent standard deviation of the mean. (**L-M**) Serum Osteopontin (OPN) and Tenascin C (TNC) concentration (ng/ml) in convalescence, stratified by radiological resolution. Statistical significance (p<0.05) tested by two way t-test. (**N-P**) Images represent data from quantitative CT analysis by Imbio (USA) lung texture analysis (LTA™) artificial intelligence based software. Images show representative LTA™ analysis results, stratified by COVID-19 convalescent disease status (Healthy control n=6; Symptomatic n=13; COVID-19 n=17). See methods for explanation of radiological terms. See Table S2 for LTA™ statistical analysis and values between groups.

TNC identified patients who developed severe COVID-19 disease, from healthy controls and mild disease (Fig.5B). TIMP1 expression significantly increased with COVID-19 disease severity, and was significantly different in mild, moderate and severe COVID-19 disease compared to baseline (Fig.5C). Further, both TNC and TIMP1 serum levels were significantly elevated in patients who died (7/30) following acute COVID-19 infection (Fig.5D-E). These data indicated our acute-injury model SOX9-panel of biomarkers have diagnostic promise in identifying patients who will develop severe acute COVID-19.

### Biomarkers downstream of SOX9 have diagnostic potential in convalescent COVID-19

During COVID recovery, patients often have persistent respiratory symptoms that are not always identified from current clinical diagnostic tests. Although persistent breathlessness may arise from multiple mechanisms, one is persistent lung injury. Recent novel imaging studies suggest conventional radiology under-estimates persistent post-COVID-19 lung injury (*39*).

To explore this further we applied automated artificial intelligence quantitative lung texture analysis™ (IMBIO, USA) to healthy control and symptomatic patient computerised tomography (CT) scans to quantify lung abnormalities across recovering groups (Table S2 and Fig.5N-P). Interstitial lung abnormalities (ground glass and reticulations) were detected in the symptomatic recovery group (9/13), despite only 6/16 patients having significant abnormalities on conventional radiology (Table S2.). To determine whether our biomarker panel could help explain these symptoms and aid diagnosis, we tested our panel in a second cohort (Fig.5F and Fig.S6) of healthy controls and recovering COVID19 patients without fibrosis. Demographics and clinical outcomes can be seen in Table S3.

TNC and OPN serum levels were increased in patients with breathlessness (Fig.5G, I) and cough (Fig. 5H, J). In contrast, lung function and radiology did not differ significantly between asymptomatic and symptomatic recovering COVID-19 patients Table S4. These results suggest TNC and OPN had diagnostic potential, identifying patients with persistent COVID-19 lung abnormalities that conventional radiology and physiology testing failed to detect, potentially signposting patients for more advanced investigations.

### Biomarkers downstream of SOX9 identify patients who have progression of lung fibrosis

Biomarkers which predict fibrotic progression for fibrotic lung disease could permit a personalised medicine approach to treatment of patients with fibrotic lung disease (*40–42*). This applies to both lung disease after COVID-19 which does not always progress, and IPF where progression is inevitable, but time to progression and speed varies between patients (*5, 6*). To address this we tested our SOX9-fibrosis panel in lung fibrosis patients (cohort 3) consisting of IPF n=12 and COVID-19 FILD n=17 (Table S3). Samples were collected at time of diagnosis and categorised based on disease progression in patients (stable, n=20; progressive, n=9) over 24 months (Fig.6A and Fig.S6A). TNC (Fig.6B), OPN (Fig.S6B) and IGFBP2 (Fig.6D) were all increased in patients with fibrotic progression compared to stable disease. Moreover, elevated TNC and IGFBP2 were prospectively associated with physiological features of deteriorating lung function (n=24; Fig.6F-H). In particular decreased forced vital capacity (FVC) and the diffusion capacity of the lungs for carbon monoxide (DLCO) trajectories, which assess lung volume and gaseous transfer across the alveolar-capillary surface associated with elevated TNC and IGFBP2 at diagnosis. As mechanistic confirmation, SOX9 was localised with TNC and IGFBP2 in human lung tissue from patients with progressive fibrosis (Fig.6C, E).

**Fig. 6.**
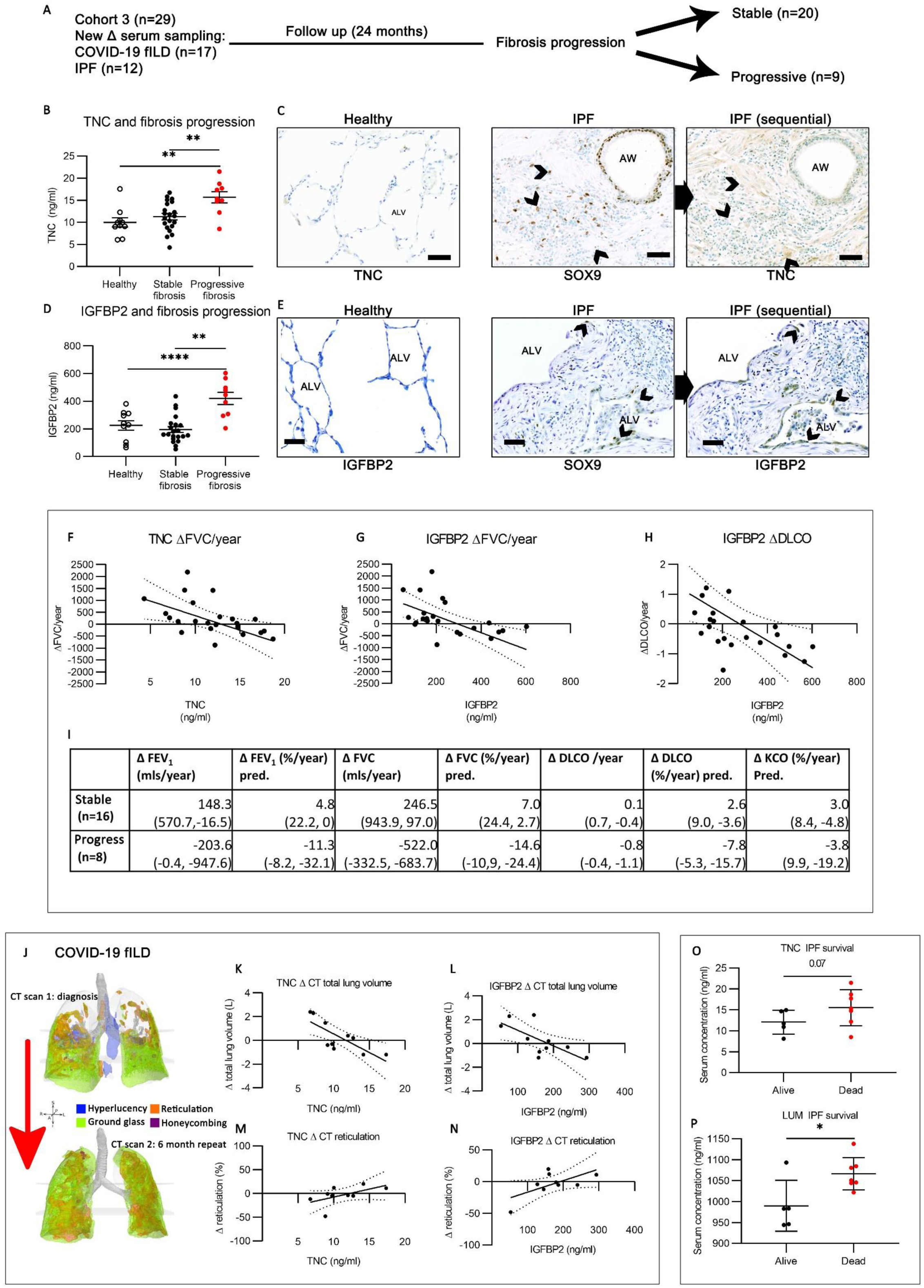
SOX9 biomarkers predict human pulmonary fibrosis progression. (**A**) Cohort 3 characteristics and outcome grouping. (**B**) TNC serum concentration (ng/ml) at diagnosis stratified by fibrosis progression. (**C**) Protein expression for TNC in healthy lung (left image). SOX9 (middle; DAB, brown) and TNC (right; DAB, brown) expression in sequential sections of human idiopathic pulmonary fibrosis lung. (**D**) IGFBP2 serum concentration (ng/ml) at diagnosis stratified by fibrosis progression. (**E**) Protein expression for IGFBP2 in healthy lung (left image). SOX9 (middle; DAB, brown) and IGFBP2 (right; DAB, brown) in the same areas (shown by black arrows). Abbreviations: AC, alveolar capillary; BV, blood vessel. Scale bars, 50μm. Counterstaining with toluidine blue. (**B**, **D**) statistical significance (p<0.05) tested by one way ANOVA. (**F**) TNC and (**G**) IGFBP2 serum concentration (ng/ml) linear regression with change in forced vital capacity (ΔFVC; ml/year), p=0.02, p=0.003. (**H**) IGFBP2 serum concentration (ng/ml) linear regression with change in predicted DLCO (ΔDLCO), p=0.01. (**i**) Change (Δ) in lung function values, per year, in stable and progressive fibrosis patients (INBUILD criteria). Units of measurement: FEV_1_, litres/min; FVC, litres; millilitres CO/minute/mm Hg. % predicted is a comparison to the GLI (2017) reference values. Values in () represent standard deviation of the mean. (**J**-**N**) Radiological change in CT scans 6 months apart in COVID-19 fILD patients, analysed using Imbio (USA) lung texture analysis (LTA™) artificial intelligence based software. (**J**) Rendered image representing quantitative LTA™ analysis of repeat scans in a COVID-19 fILD patient. (**K**) TNC and (**L**) IGFBP2 linear regression with Δtotal lung volume (L/year), p=0.009, p=0.03. (**M**) TNC (ng/ml) linear regression with Δreticulations (%), p=0.12. (**N**) IGFBP2 (ng/ml) linear regression with Δreticulations (%), p=0.07. (**O**) TNC (p=0.07) and (**P**) Lumican (LUM; p=0.01) serum concentration (ng/ml) at diagnosis, stratified by 3 year survival status. Statistical significance (p<0.05) determined by two way t-test.

We next assessed the capability of our SOX9-panel to predict disease-specific measures of progression. In a subset of our COVID-19 FILD patients (10/18), CT scans had been carried out at diagnosis and 6 months follow up (Fig.6J). We applied quantitative lung texture analysis™ (IMBIO, USA) to compare and quantify fibrotic radiological changes (see methods). Increased TNC and IGFBP-2 (Fig.6K, L) were significantly associated with reduced total lung volume. Moreover, TNC, IGFBP-2 and OPN were all associated with change in CT ground glass (Fig.S7F-H) and lung reticulation, which represent pathological changes to lung interstitium (Fig.6M, N and Fig.S7K). In our IPF patient cohort, LUM and TNC were elevated in serum from IPF patients that died within the following 3 years (Fig.6O, P) No patients with COVID-19 FILD died within the follow-up period.

Collectively these results show that our SOX9-panel has the potential to identify patients who will develop clinically meaningful adverse physiological, radiological and mortality outcomes in human lung fibrosis.

## Discussion

SOX9 has an established importance to alveolar regenerative populations, with early phase human work already underway to therapeutically target these cells (*28*). Most *in vivo* lung studies, to date, of SOX9 have focussed upon this role of SOX9 (*43, 44*). Our results here indicate SOX9 has an importance to lung injury and fibrosis reaching beyond this. We show here that SOX9 is critical to dysregulated alveolar epithelial and fibroblast injury responses, two critical components of the fibrotic niche. *Sox9 KO* subsequently protects against severe acute lung injury and pulmonary fibrosis.

Our, and other groups, previous work has established that SOX9 directly transcriptionally regulates a number of ECM proteins in organ development and fibrosis (*10, 34, 45, 46*). To date, the only *in vivo* study of SOX9 addressing progressive pulmonary fibrosis uses Col1⍺2-Cre^ER^ (fibroblast-specific) conditional knockout and over-expression models and is consistent with our work in other organs (*10, 34, 45, 46*). Here, our *in vivo* work supports SOX9 regulating ECM secretion in lung fibroblasts in pulmonary fibrosis. However, the molecular role of SOX9 in fibrotic transitional epithelial (KRT17+ or SPRR1A+) cells and acute lung injury is currently unknown. Our *in vivo* data shows KRT17 and SPRR1A are expressed in damage associated alveolar epithelium from an early time-point in lung injury, and *Sox9* loss reduced the expression of these cells in alveoli. Such a role is consistent with developmental organoid work, showing SOX9 maintains a proliferating alveolar progenitor population but prevents terminal differentiation (*23*).

Determining whether SOX9 regulates damage associated transitional epithelium due to direct effects of SOX9 on gene transcription within these cells, or indirectly through mediating cross-talk within the fibrotic niche, may provide important insights into how this disease progresses. The *in vivo* importance of SOX9 to alveolar epithelial damage responses and pro-fibrotic fibroblasts in lung injury, identified in our work, has clear translational significance to COVID-19 and IPF where dysregulated alveolar damage responses and excessive fibroblast ECM secretion drives severity in both conditions (*47, 48*). Thus, targeting molecular pathways up- and down-stream of SOX9 in transitional epithelial cells and ECM secreting fibroblasts may provide sorely needed novel therapeutic targets.

Facilitating early diagnosis and treatment of both severe acute COVID-19 lung injury and progressive lung fibrosis is vital to improving outcomes and reducing long-term morbidity of patients. In determining the *in vivo* role of SOX9 we identified secreted ECM biomarkers downstream of SOX9 in our *Sox9 KO* models. In hospitalised acute COVID-19 and lung fibrosis patient serum, selected biomarkers showed ability in identifying patients who will develop adverse changes to lung physiology (maximal FiO_2_, FVC and DLCO) and important, clinically meaningful outcomes (COVID-19 severity, progression of fibrosis and mortality). These data were reflective of the pathological role of SOX9 shown in our *in vivo* models. Clearly biomarkers identified here need to be validated in larger prospective cohorts. However, if replicated, our SOX9-panel of biomarkers have the potential to guide enhanced monitoring and earlier treatment of high-risk hospitalised COVID-19 patients, as well as earlier diagnosis and antifibrotic treatment in PFILD patients.

This study shows that SOX9 is functionally critical to disease in acute lung injury and pulmonary fibrosis. It has translational importance with diagnostic, prognostic and therapeutic potential to both COVID-19 and IPF.

## Methods

### Animal experiments

RosaCreER:*Sox9*^fl/fl^ animal experiments were carried out with approval from the University of Manchester Animal Welfare and Ethical Review Board and in accordance with UK Home Office regulations. KRT18-hACE2 experiments were carried out by Professor Hanbro’s laboratory group (Centenary Institute for Inflammation, University Technology Sydney, Australia), as previously published (*29*) with approval from the Sydney Local Health District (SLHD) Animal Ethics and Welfare Committee and adhering to the Australian Code for the Care and Use of Animals for Scientific Purposes (2013) as set out by the National Health and Medical Research Council of Australia. SARS-CoV-2 mouse infection experiments were approved by the SLHD Institutional Biosafety Committee (*29*). All experimental mice were on a C57BL/6 J background in housing with a 12-hour light dark cycle and food and water available *ad libitum* (*29, 33*).

RosaCreER:*Sox9*^fl/fl^ experiments: RosaCreER:*Sox9*^fl/fl^ animals have been described previously (*33*). Briefly, tamoxifen (100 mg Kg^-1^; Sigma, UK) was injected i.p. to activate CreER activity and induce *Sox9* deletion in ROSACreER:*Sox9*^fl/fl^ animals. ROSACreER^+/−^ and ROSACreER^−/−^ animals were injected with tamoxifen to control for any unexpected effects. For acute lung injury *in vivo* experiments tamoxifen was injected 8, 7 and 5 days prior to bleomycin treatment, with lungs harvested 3 days post-bleomycin. For fibrosis *in vivo* experiments tamoxifen was injected 5, 6 and 8 days after bleomycin administration with lungs harvested at day 14 post-bleomycin. Bleomycin (0.003IU/Kg; Kyowa-Kirin, Japan) was administered via the oropharyngeal route (*49*).

Mice were exsanguinated, prior to lavage of bronchoalveolar space with 0.8 ml 2mM EDTA PBS solution (4°C) using a blunted 20-gauge needle inserted into the trachea. BALF was centrifuged at 450g for 5 minutes, with BALF supernatant snap frozen and stored at -80°C. Lungs were then inflated with 4% paraformaldehyde solution and tied off, prior to paraffin embedding and sectioning (5μm) for IHC experiments. In separate experiments lungs were dissected deflated and snap frozen.

KRT18-hACE2 experiments: The KRT18-hACE2 SARS-CoV-2 lung injury model has been published previously (*29*). Mice were anaesthetised with isoflurane followed by intranasal challenge with 10^3^ PFU SARS-CoV-2 (VIC01/2020) in a 30 µL volume (*29*). Infection and injury were confirmed using lung and BALF viral titre, BALF absolute immune cell count and histology as published previously (*29*). At day 6 post-infection, mice were euthanised with intraperitoneal overdose of pentobarbitone (Virbac, Australia) (*29*). The single lobe lung was perfused with 0.9% NaCl solution via the heart, followed by inflation with 0.5 mL 10% neutral buffered formalin through the trachea, and storage in 10% neutral buffered formalin (*29*). Following fixation for at least 2 weeks, single lobes were transported to a PC2 facility where they were paraffin-embedded, sections cut to 3 µm thickness using a Leica microtome (Leica, Germany), then samples were transferred to the Piper Hanley laboratory, University of Manchester for IHC experiments.

Hydroxyproline assay was performed as previously published, (*50*) with some established modifications (*51*). Whole lungs were harvested, with extrapulmonary vessels and airways excised and weighed. Lungs were flash frozen and powdered prior to drying in an oven at 80°C for 1 hour. Dried lung was hydrolysed for 20 hours in 6N HCl. Hydrolysed sample (20μl) was added to (100μl) chloramine T (Sigma, UK) solution prepared in a citrate-acetate buffer and incubated for 20 minutes at room temperature. Subsequently Erlich’s solution was prepared (4-dimethylaminobenzaldehyde, Sigma, UK) added and incubated at 60°C for 30 minutes. After this colorization the plate was read using the Glomax multi microplate reader (Promega) at an absorbance wavelength of 560nm. Standard curves were generated for each experiment using reagent hydroxyproline (Merck) as a standard. Results were expressed as micrograms of hydroxyproline contained per each lung total tissue.

Western blot protein expression was analysed as previously described (*33, 52*). Total protein was used as loading control. Quantity One software (Bio-Rad) was used for image acquisition and data analysis. Antibodies used for western blot can be found in Table S8.

Flow cytometry: BALF was collected and the cellular fraction isolated as described above. Pelleted cells were then re-suspended, manually counted using tryptan blue and haemocytometer prior to immunostaining. Flow cytometry antibodies can be seen in Table S7. Flow cytometric measurements were performed using the Fortessa LSRII (BD biosciences) and FACS DIVA™ software (BD biosciences). Results were analysed with FlowJO software (BD biosciences). Compensation was performed using UltraComp eBeads (eBioscience). Cells passed through the cytometer were gated by granularity and size to select single cells. Live/Dead Blue (Invitrogen) was used to select live cells and fluorescence minus one (FMO) control staining was used. CD45^+^ cells were gated as shown in Fig.S5.

Proteomic analysis of bronchoalveolar lavage: BALF supernatant was collected and isolated as described above. In brief supernatant protein concentration was standardised prior to being concentrated using Microcon® (Merck) 10Kda filter columns and centrifugation (14,000g 20 minutes). Samples were reduced and alkylated prior to S-Trap™ plate and trypsin peptide digestion. After desalting and cleaning protein was vacuum dried to completeness (Heta Vacuum centrifuge, Thermofisher). Samples were analysed using LC-MS on the Q Exactive™ HF (Thermofisher) mass spectrometer.

Unbiased whole proteome analysis was performed, according to treatment (bleomycin or saline) and genetics (*WT* or *Sox9 KO*) with acute lung injury and pulmonary fibrosis experiments analysed separately (Fig.4A and E). Differentially expressed proteins were organised into κ-means clusters (Fig.4A and E), which were annotated using gene ontology (GO) analysis (Fig.4B and F), and defined by a fold-change >log2 (1) and p<0.05.

### Human samples

Cohort 1: Patients were admitted with acute COVID-19 pneumonitis between April 2020 and December 2020 across two hospitals within Manchester Foundation NHS Trust. Diagnosis of COVID-19 was either proven with polymerase chain reaction (PCR) testing for SARS-CoV-2 or considered likely according to clinical judgement of specialist respiratory physicians, based upon radiological changes seen on chest radiographs or computed tomography (CT) imaging, consistent with COVID-19 pneumonitis, combined with typical signs or symptoms of COVID-19. Patients were excluded if COVID-19 was not their primary diagnosis. Serum samples were taken at time of inpatient admission. Inpatient severity was defined by their maximal respiratory support requirement to maintain SpO_2_ >90% as: mild, FiO_2_ <0.28; moderate, ≤ 0.28 FiO_2_ <0.6; Severe, FiO_2_ ≥0.6, or non-invasive mechanical ventilation, or intubation and mechanical ventilation. These severity criteria were designed to be inclusive of patients not suitable for invasive monitoring or intubation and mechanical ventilation due to either frailty, comorbidity or personal choice and matched hospital treatment guidelines at the time.

Cohort 2: Patients were recruited at their 1^st^ outpatient appointment (3-6 months) after being hospitalised with COVID-19 pneumonitis two hospitals within Manchester Foundation NHS Trust (MFT). Patients met the same inpatient criteria as cohort 1. Fibrosis was excluded using radiology (CXR available in 25/26; CT scan available in 13/26), lung function (available 23/26) and clinical expertise of specialist respiratory physicians.

Cohort 3: Patients with COVID-19 FILD or IPF were recruited at time of diagnosis in tertiary ILD outpatient clinics at Wythenshawe Hospital, MFT. All patients with lung fibrosis (COVID-19 FILD, or IPF) included in the study had their diagnosis confirmed by a tertiary interstitial lung unit multi-disciplinary team that includes a specialist thoracic radiologist, a thoracic histopathologist, a lung transplant physician and specialist interstitial lung disease physicians. All patients (29/29) had high resolution CT scans and lung function physiological testing at time of diagnosis. Patients were prospectively followed up according to usual standard of care for their disease (including repeat radiology and lung function physiological tests). Progression of fibrosis was determined according to INBUILD progressive fibrosing ILD trial criteria, over 24 months.(*42*) Patient survival was followed up for 3 years in patients with IPF.

Quantitative CT analysis: CT thorax scan images from convalescent COVID-19 patients (fibrotic 17/17 and non-fibrotic 13/26) taken at follow up appointments at MFT. Repeat HRCT scans performed 6 months after initial scan were available in 10/17 COVID-19 fILD patients. CT scans were analysed using Lung Density Analysis™ and Lung Texture Analysis™ programmes (Imbio Inc., Minneapolis, MN, USA) to generate quantitative CT reports (ManARTS biobank (15/NW/0409); study number M2022-119). This generated quantitative scan reports, as shown in Fig. 5N-P and Fig.6J. Hyperlucency is a radiological abnormality that can represent alveolar, bronchiolar or vascular damage. Ground glass and reticular radiological abnormalities represent interstitial changes which can be seen in fibrotic lung disease as well as other pathologies. Honeycombing represents architectural destruction in regions of advanced fibrosis. Total lung volume reduces with fibrosis. Pulmonary vessel volume (PVV) increases with fibrosis. Further information on this technology can be found in the Imbio LDA+ user manual version 4.1.0 and the lung texture analysis user manual version 2.2.0. https://www.imbio.com/support-documentation/Serum samples were collected at time of diagnosis in each of the three cohorts and stored at -80°C (ManARTS study numbers M2020-93, M2014-18).

Serum protein quantification was performed using Luminex multi-plex assays (R&D systems), which utilise magnetic beads coated with antibodies specific to analytes (TIMP1, OPN, TNC, IGFBP2, LUM). Measurements were performed by the Bio-Plex 200 system (Bio-Rad). Accuracy was measured with coefficients of variation for each marker assayed and a sample was discounted if variation was >10% between replicates.

Lung tissue was collected from ex-plants of patients receiving transplant for idiopathic pulmonary fibrosis, with samples formalin fixed and paraffin embedded (REC 14/NW/0260). Control tissue was from healthy margins of patients with resections for high-risk lung nodules (ManARTS study number M2014-18).

Immunohistochemistry: Human and animal tissue sections were prepared as described, and IHC performed as previously published (*33, 34*). Briefly, heat-induced epitope retrieval was performed in sodium citrate buffer (pH 6). Detection was performed using primary antibodies listed in Table S5 followed by incubation with either anti-rabbit ImmPRESS AP reagent and development with ImmPACT Vector Red substrate (Vector), or incubations with species-specific biotinylated antibodies (Table S6; Vector) and then streptavidin–horseradish peroxidase (HRP) antibody (Vector; 1:200) before development with diaminobenzidine (Sigma-Aldrich). Counterstaining was performed using toluidine blue. PSR staining was performed as previously described (*33, 52*).

## Acknowledgements

The Platform Technologies at the University of Manchester are acknowledged for their technical help and support. This report is independent research supported by the North West Lung Centre Charity at Manchester University NHS Foundation Trust. The authors would like to acknowledge the Manchester Allergy, Respiratory and Thoracic Surgery Biobank and the North West Lung Centre Charity for supporting this project. The views expressed in this publication are those of the author(s) and not necessarily those of the NHS, the North West Lung Centre Charity or the Department of Health.

## Author Contributions

KPH, LP and NAH conceived and designed experiments. LP and EJ contributed to experimental planning and design. JFB, PH, AS, RS, RV, SS, CH and PR-O provided reagents and contributed to experimental design. LP, EJ, KS, LB, LM, and NS performed experiments. LP, EJ and KPH analysed the data. YO and CL carried out bioinformatics and statistical analysis. KPH and LP guided experiments and analysed data. KPH, JFB, LP and EJ wrote and edited the manuscript.

## Data and Materials Availability

The proteomic data will be deposited into (XXXX; added on acceptance) with the identifier (XXXX; added on acceptance). All other data needed to evaluate the conclusions in the paper are present in the paper or the Supplementary Materials.

---

**Fig. S1.**
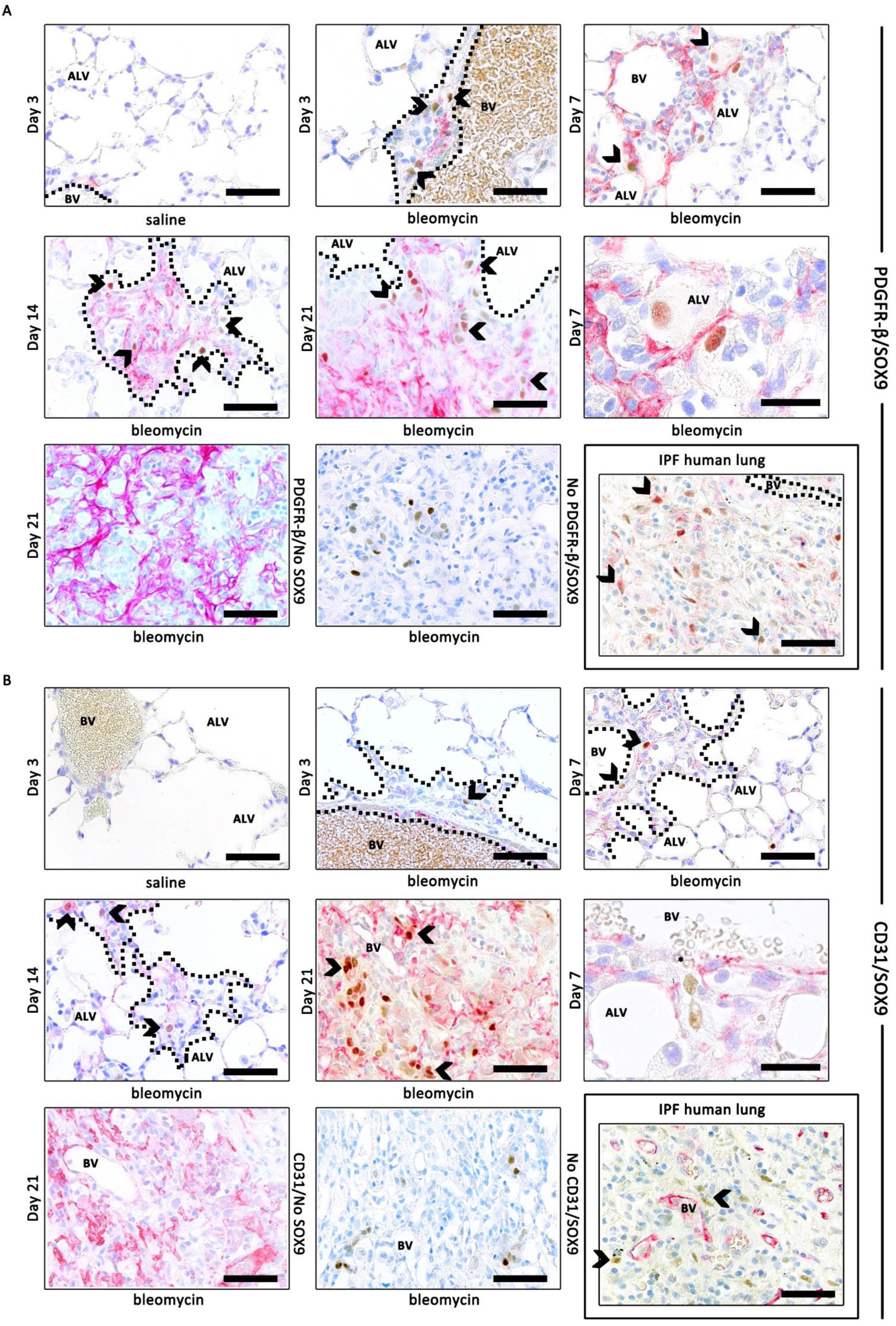
Co-localisation of SOX9 with stroma (PDGRβ+ cells) and vasculature (CD31+ cells) throughout the wild type bleomycin injury time-course. (A) Top left shows SOX9/PDGFRβ expression in control lung. SOX9 (DAB, brown) and PDFGR-β (red) expression is shown in injured lung throughout the time-course. Regions of architectural destruction by stroma are identified by black dotted lines. Middle right image shows SOX9/PDGFR-β expression within injured alveoli at high magnification (day 7). Bottom right image shows SOX9 (DAB, brown) and PDGFRβ expression in human IPF lung. (**B**) Top left shows SOX9 (DAB, brown) CD31 (red) expression in control lung. Subsequent images show SOX9 (DAB, brown) and CD31 (red) expression in bleomycin injured lung throughout the time-course. Middle right image shows SOX9/CD31 expression proximal to damaged endothelium, of a blood vessel (BV) at high magnification. Bottom right image shows SOX9 (DAB, brown) and CD31 (red) expression in human IPF lung. Single staining controls can be seen in the bottom row (**A**, **B**). Scale bars, 50 μm aside from high magnification (20 μm). Counterstaining toluidine blue. Abbreviations: ALV (alveoli); BV (blood vessel).

**Fig. S2.**
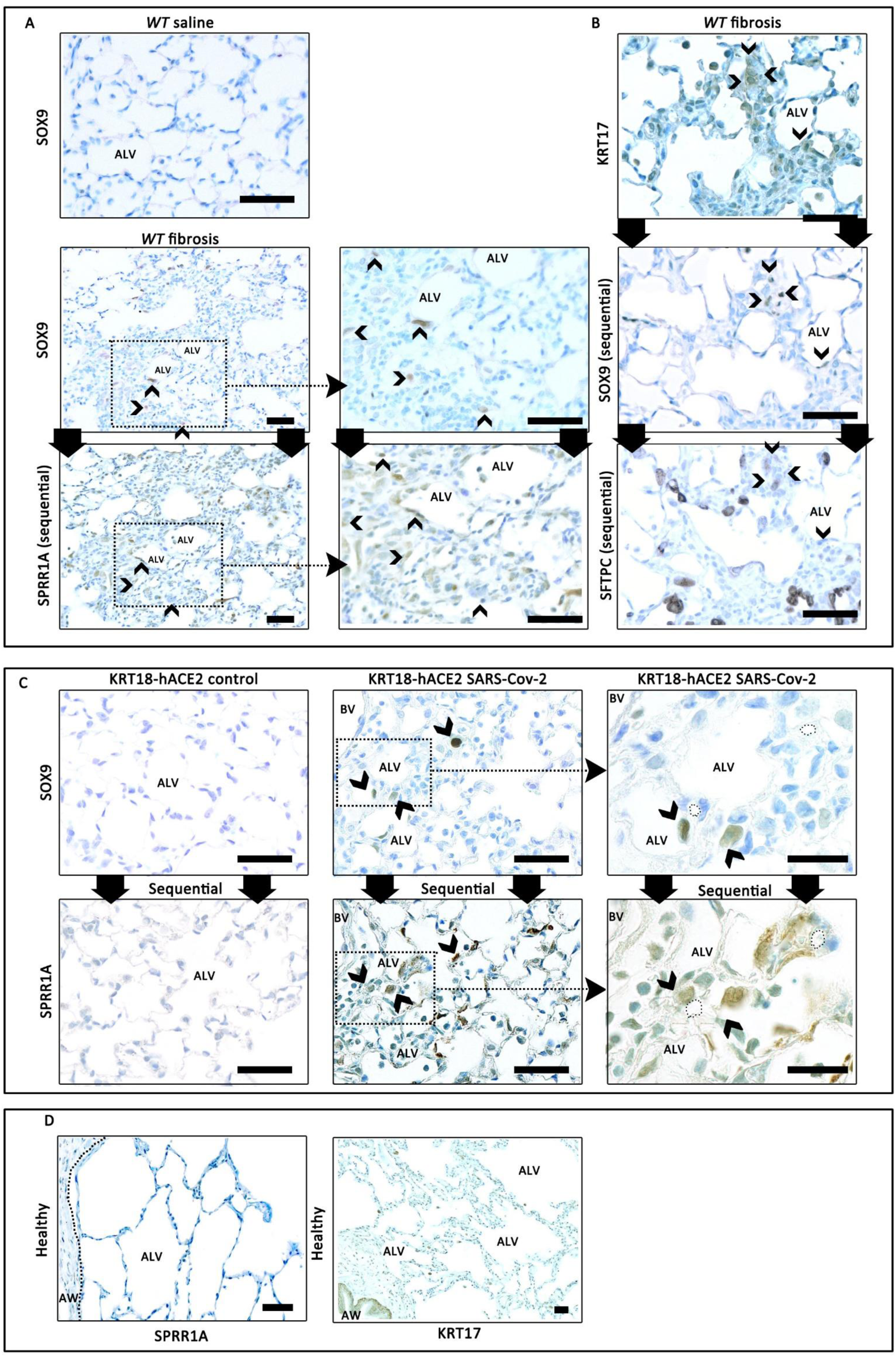
SOX9 co-localises to damage associated alveolar epithelial markers SPRR1A and KRT17 in *Sox9 KO* fibrosis and KRT18-hACE2 SARS-CoV-2 models. (**A**, **B**) Experiments from *Sox9 KO* fibrosis model animals. (**A**) Top image shows SOX9 expression in *wild type (WT)* saline control lung. Middle image shows SOX9 expression in *WT* bleomycin fibrotic lung, bottom image shows SPRR1A expression (both DAB, brown) in sequential section. Right hand image shows magnified area (dotted black line). (**B**) KRT17 (top), SOX9 (middle) and pro-SFTPC (bottom) expression in sequential section of *WT* bleomycin fibrotic lung. (**C**) Experiments from KRT18-hACE2 SARS-CoV-2 model animals. Top left shows SOX9 and SPRR1A (DAB, brown) in sequential sections of control lung. Middle shows SOX9 and SPRR1A expression (DAB, brown) of SARS-CoV-2 injured lung in sequential sections. Right hand images show magnified areas (dotted black line). (**D**) SPRR1A and KRT17 expression in healthy control human lung. Expression in fibrotic human lung can be seen in Fig. 2I. Counterstain toluidine blue. Scale bars, 50 μm, apart from high magnification images (20 μm).

**Fig. S3.**
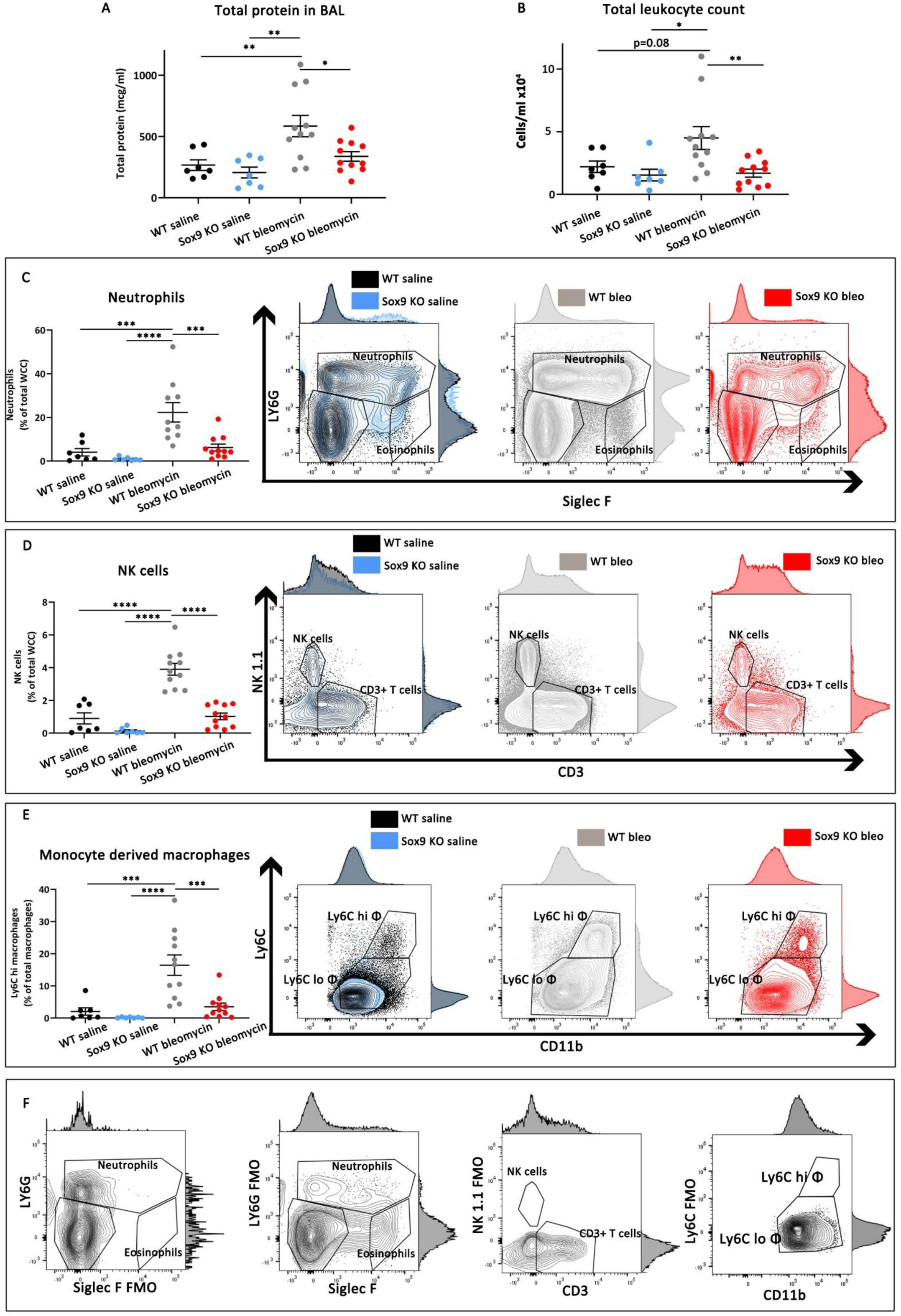
*Sox9 knockout (Sox9 KO)* prior to injury protects against acute lung injury severity and reduces immune cell recruitment to the lung. All images represent experiments performed on bronchoavleolar lavage (BAL) fluid from *wild type (WT)* and *Sox9 knockout (Sox9 KO)* acute lung injury model animals (3 days; n=7-11 per group). (**A**) Bronchoalveolar (BAL) total protein content standardised for retrieval volume (μg/ml). (**B**) Total leukocyte (TER119^-^CD45^+^) cell numbers, standardised for BAL retrieval volume (cells/ml). (**C**) Neutrophils (TER119^-^CD45^+^CD64^-^MERTK^-^LY6G^+^) total leukocyte proportion (%WCC). (**D**) Natural killer (NK) cell (TER119^-^CD45^+^CD64^-^MERTK^-^Ly6G^-^SiglecF^-^CD3^-^NK1.1^+^) total leukocyte proportion (%WCC). (**E**) Proportion of total macrophage population (TER119^-^ CD45^+^CD64^+^MERTK^+^) derived from monocytes (TER119^-^CD45^+^CD64^+^MERTK^+^Ly6C^hi^). (**F**) Fluorescence minus one (FMO) control staining and population gates. Statistical significance (p<0.05) tested by two-way ANOVA.

**Fig. S4.**
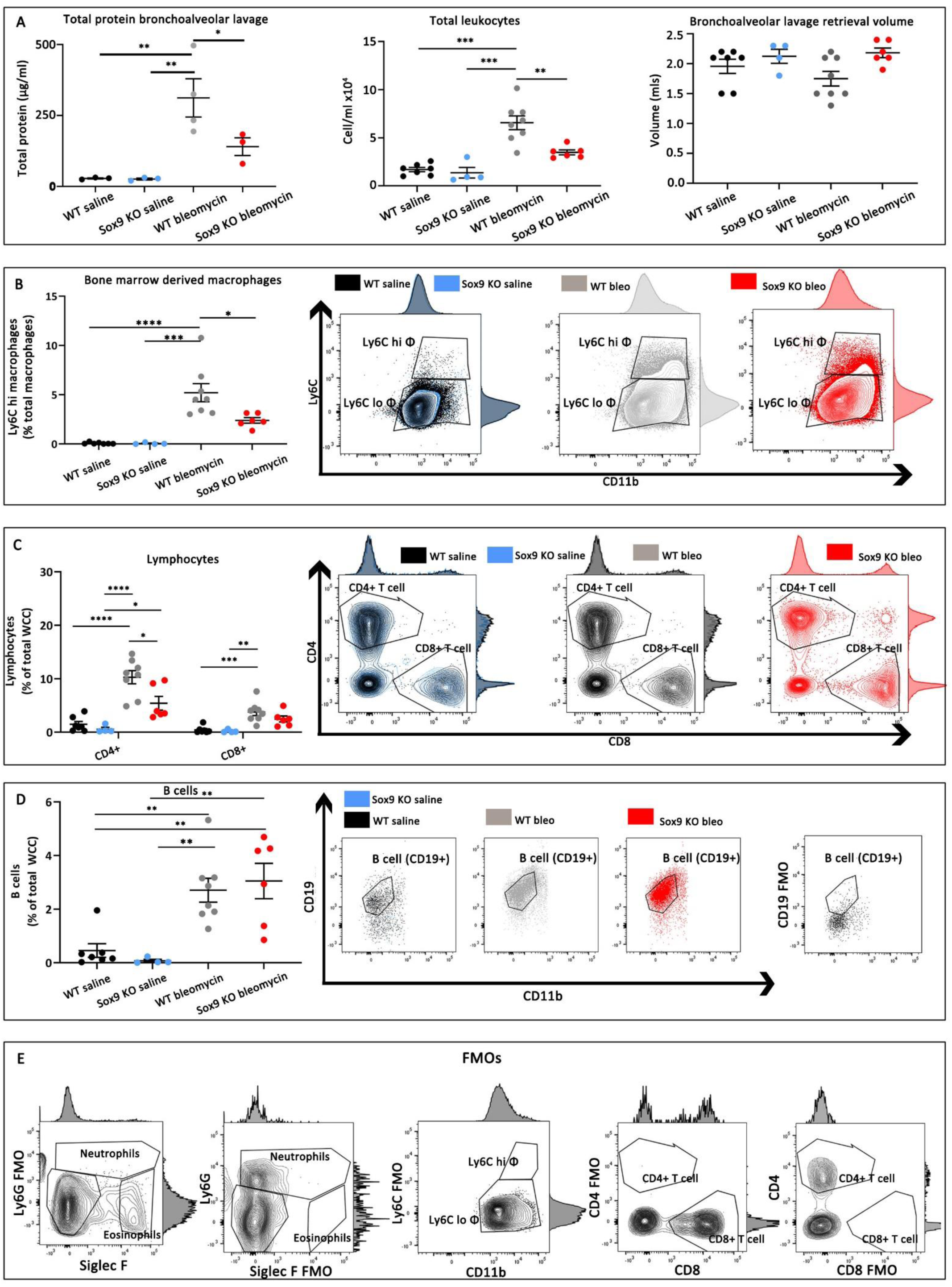
*Sox9 knockout (KO)* during fibroproliferation reduces immune cell recruitment to the lung. All images represent experiments performed on bronchoavleolar lavage (BAL) fluid from *wild type (WT)* and *Sox9 knockout (Sox9 KO)* lung fibrosis model animals (14 days; n=4-8 per group). (**A**) Left graph shows bronchoalveolar (BAL) total protein content standardised for retrieval volume (μg/ml). Middle graph shows total leukocyte (TER119^-^CD45^+^) cell numbers, standardised for BAL retrieval volume (cells/ml). Right graph shows retrieval volume. (**B**) Proportion of total macrophage population (TER119^-^CD45^+^CD64^+^MERTK^+^) derived from monocytes (TER119^-^CD45^+^CD64^+^MERTK^+^Ly6C^hi^). (**C**) CD4^+^ and CD8^+^ lymphocyte (TER119^-^CD45^+^CD64^-^MERTK^-^LY6G^-^SiglecF^-^MHCII^-^CD11b^-^NK1.1^-^CD3+) proportion of total leukocyte population (TER119^-^CD45^+^; %WCC). (**D**) B cell (TER119^-^CD45^+^CD64^-^ MERTK^-^LY6G^-^SiglecF^-^CD11b^-^MHCII^+^CD19^+^) proportion of total leukocytes (TER119^-^ CD45^+^; %WCC). (**E**) Fluorescence minus one (FMO) control staining and population gates. Statistical significance (p<0.05) tested by two-way ANOVA.

**Fig. S5.**
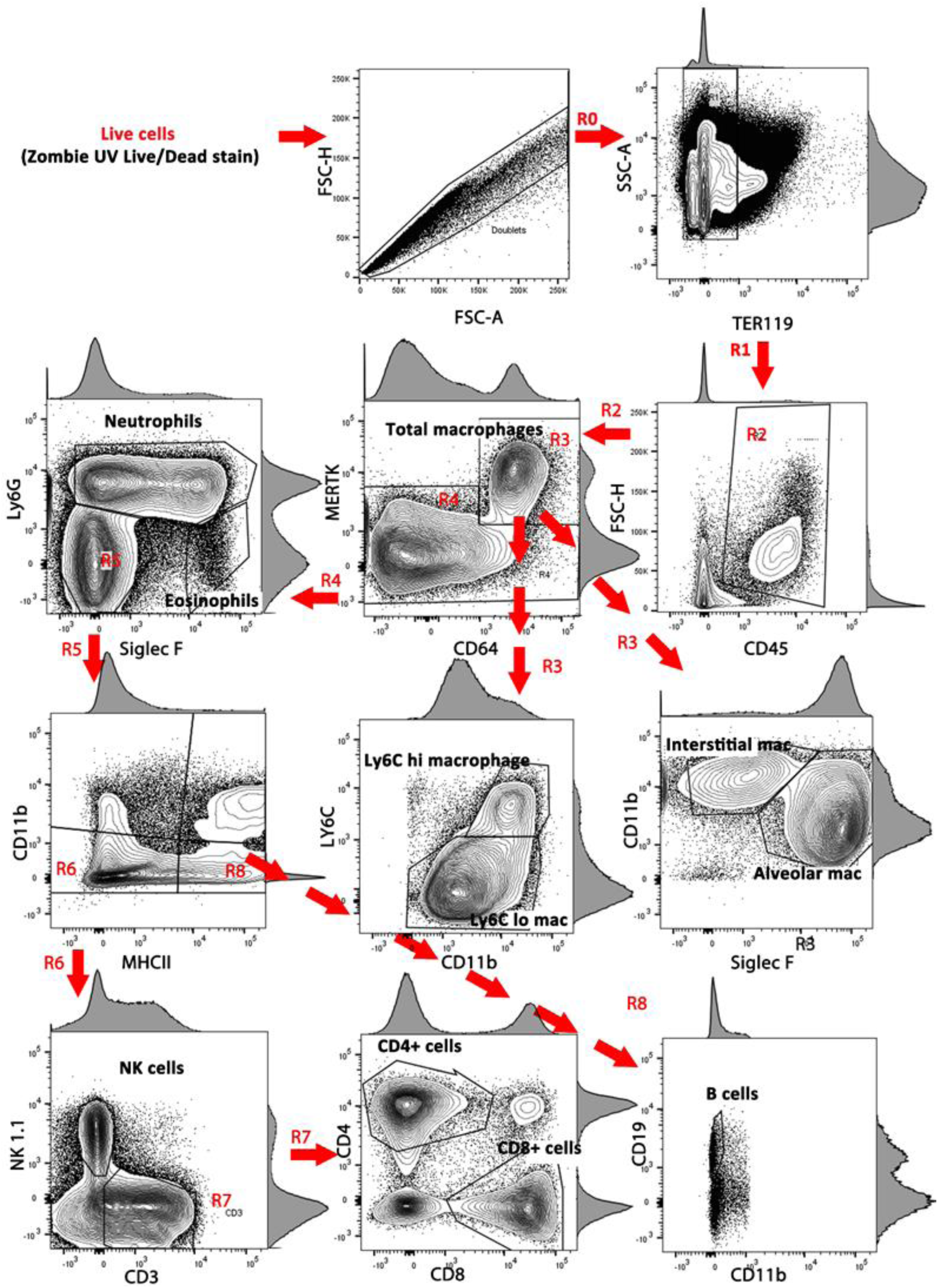
Flow cytometry gating strategy. Gating strategy of flow cytometry analysis for experiments shown in Fig. S3 and S4. Frequency histograms shown on log transformed X and Y axes.

**Fig. S6.**
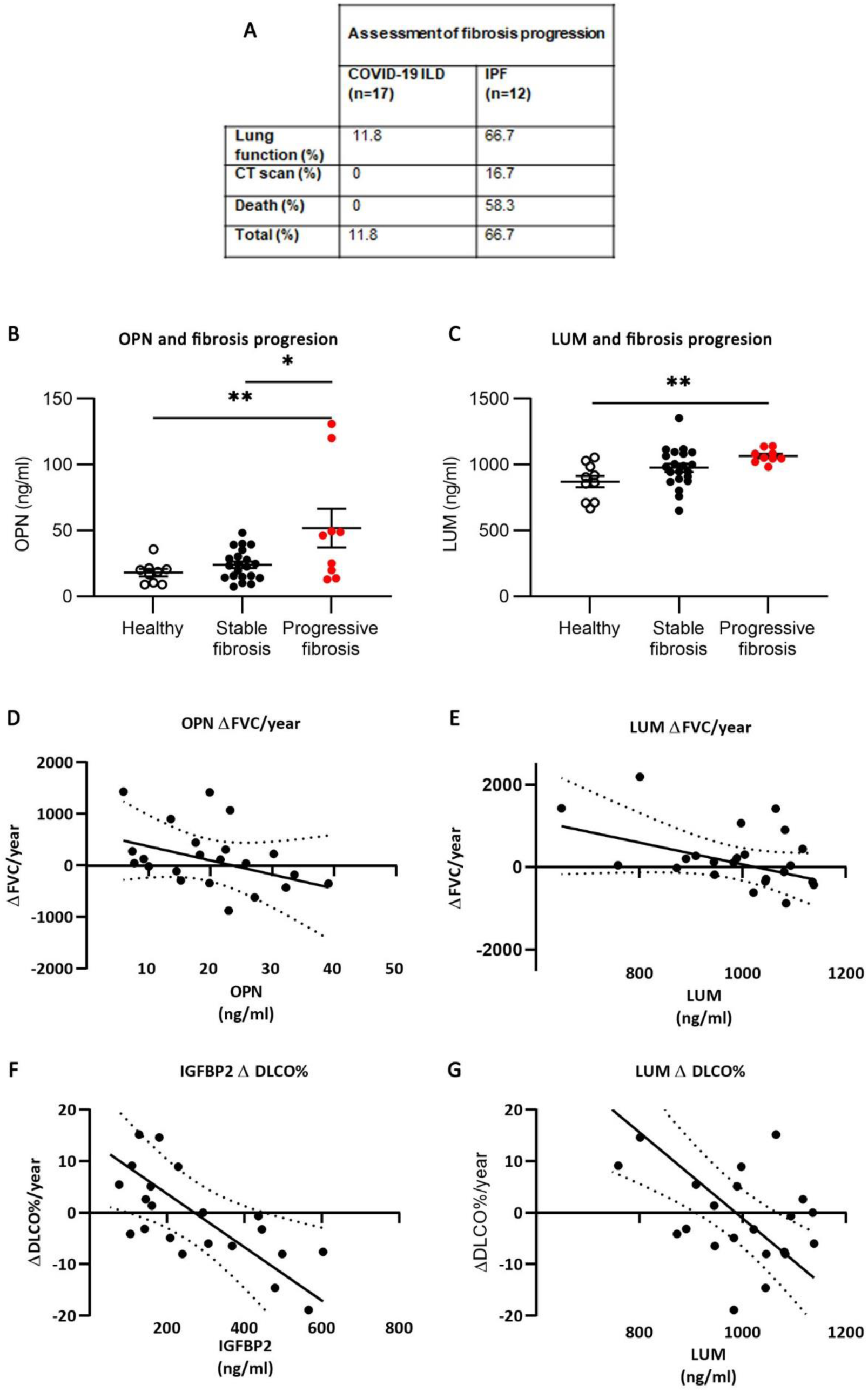
Additional fibrosis progression biomarker data. (**A**) Table showing method of diagnosing fibrosis progression in COVID-19 fibrosing interstitital lung disease (FILD; n=17) and idiopathic pulmonary fibrosis (n=12). Note patients may have more than one measure identifying progression. (**B**) Osteopontin (OPN) and (**C**) LUM serum concentration at time of diagnosis stratified by subsequent fibrosis progression (2 years). Statistical significance (p<0.05) tested by one way ANOVA. (**D-G**) Scatter plots showing linear regression of serum biomarker (ng/ml) with future Δ lung function. OPN (**D**) and LUM (**E**) did not have significant association (p>0.05) with ΔFVC (mls/year). (**I**) IGFBP2 (p=0.01) and (**J**) LUM (p=0.001) predicted ΔDLCO(%). Dotted lines represent 95% confidence intervals. Units of measurement: FEV_1_, litres/min; FVC, litres; millilitres CO/minute/mm Hg. % predicted is a comparison to the GLI (2017) reference values. Values in () represent standard deviation of the mean.

**Fig. S7.**
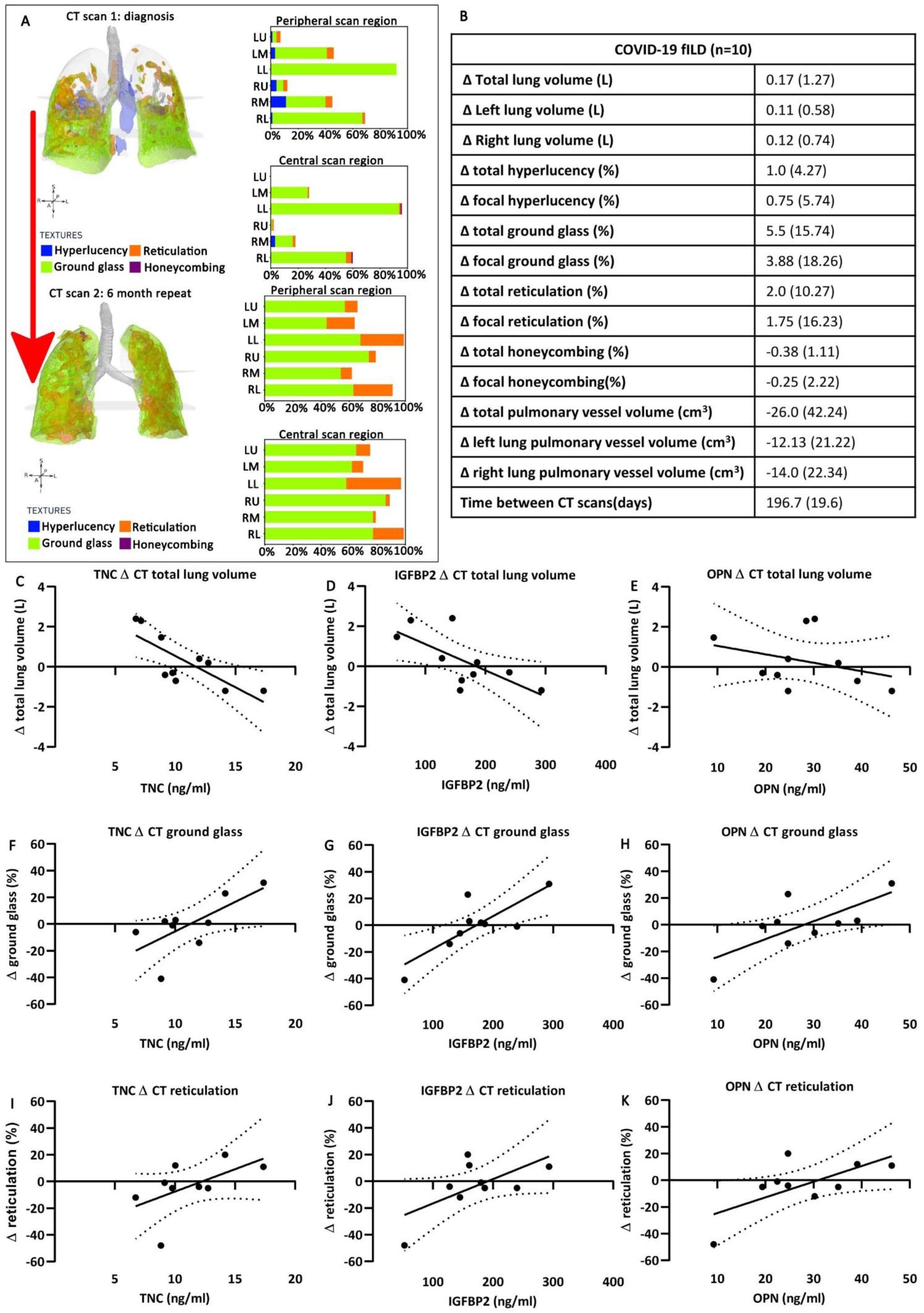
Biomarkers predict radiological progression in COVID-19 fibrosing interstitial lung disease (FILD). Radiological change in CT scans 6 months apart in COVID-19 FILD patients, analysed using Imbio (USA) lung texture analysis (LTA™) artificial intelligence based software (n=10). (**A**) Rendered image representing quantitative LTA™ analysis of repeat scans in a COVID-19 FILD patient. and quantitative output (% affected regions). (**B**) Table showing change (Δ) in LTA™ quantitative CT analysis values. (**C-K**) scatter plots showing diagnosis serum biomarker expression (ng/ml) regression (line) with ΔLTA™ quantitative CT analysis values. (**c**) TNC predicts Δtotal lung volume (L/year), p=0.009. (**D**) IGFBP2 predicts Δtotal lung volume (L/year), p=0.03. (**E**) OPN does not predict Δtotal lung volume (L/year), p=0.35. (**F**) TNC predicts Δground glass (%), p=0.04. (**G**) IGFBP2 predicts Δground glass (%), p=0.009. (**H**) OPN predicts Δground glass (%), p=0.03. (**I**) TNC does not predict Δreticulations (%), p=0.12. (**J**) IGFBP2 does not predict Δreticulations (%), p=0.07. (**k**) OPN predicts Δreticulations (%), p=0.05. Abbreviations: LU, left upper lung; LM, left middle lung; LL, left lower lung; RU, right upper lung; RM, right middle lung; RL, right lower lung; TNC, tenascin C; IGFBP2, insulin growth factor binding protein 2; OPN, osteopontin. Values in () represent standard deviation of the mean. Dotted lines represent 95% confidence intervals.

**Table S1:** Cohort 1 (acute hospitalised COVID-19) patient characteristics. Table showing acute COVID-19 patient (cohort 1) demographics, co-morbidities and outcomes stratified by severity. Severity was defined as: mild, maximal FiO_2_≤0.24; moderate, maximal FiO_2_>0.24 <0.6; severe, maximal FiO_2_≥0.6 or requiring acute non-invasive ventilation or invasive mechanical ventilation. Statistical significance (p<0.05) was determined by one way ANOVA. One patient died within 30 days from a complication of COVID-19 infection, all others due to pneumonitis.

|  | All<br>n=30 | Mild<br>n=9 | Moderate<br>N=12 | Severe<br>n=9 | p value<br>(**<0.05**) |
| --- | --- | --- | --- | --- | --- |
| **Age** | 61.0<br>(16.6) | 61.3<br>(20.1.) | 61.2<br>(15.0) | 64.3<br>(11.9) | NS |
| **Sex**<br>**(M:F)** | 1.58 | 2.0 | 0.5 | 8.0 | p< 0.05 |
| **Ethnicity**<br>**(Caucasian %)** | 77.4 % | 67.7 % | 75.0 % | 100 % | NS |
| **BMI** | 29.7<br>(8.9) | 26.3<br>(9.2) | 32.4<br>(10.0) | 31.2<br>(4.1) | NS |
| **Hypertension** | 67.7 % | 66.7 % | 58.3 % | 77.8 % | NS |
| **IHD** | 19.4 % | 22.2 % | 16.7 % | 22.2 % | NS |
| **Pulmonary<sup>†</sup>** | 12.9 % | 22.2 % | 8.3 % | 11.1 % | NS |
| **Asthma** | 16.1 % | 11.1 % | 33.3 % | 0.0 % | p<0.01 |
| **Diabetes** | 32.2 % | 0.0 % | 33.3 % | 66.6 % | 0.03 |
| **CRP** | 113.7<br>(110.5) | 34.4<br>(31.7) | 89.0<br>(53.0) | 244.4<br>(106.9) | <0.001 |
| **LOS**<br>**(days)** | 15.5<br>(14.7) | 11.3<br>(14.7) | 11.4<br>(9.7) | 26.0<br>(13.9) | 0.002 |
| **30 day mortality**<br>**(deceased/alive)** | 19.4 % | 0.0 % | 8.3 % | 66.6% | <0.001 |

**Table S2:**
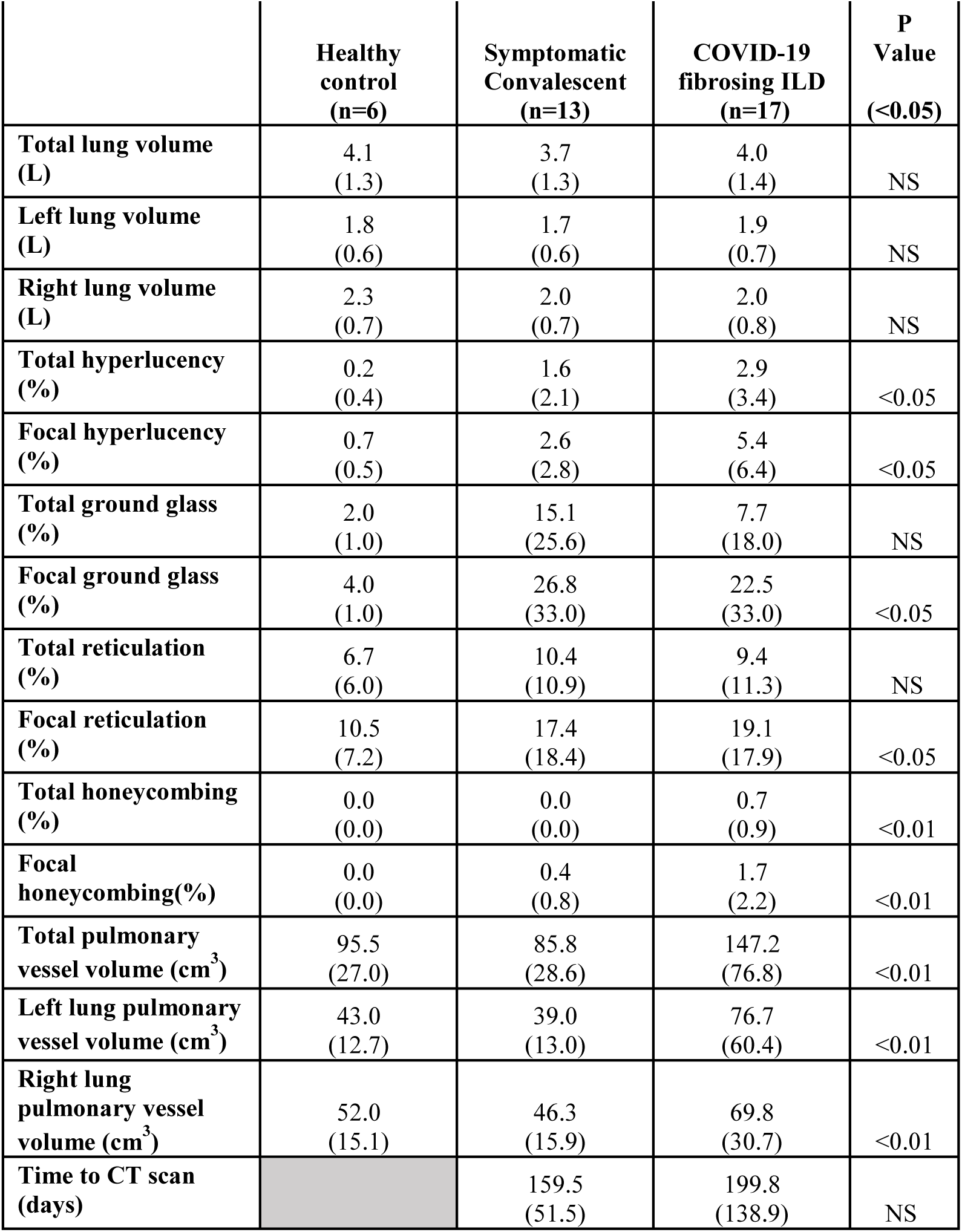
Quantitative CT scan analysis shows abnormalities in symptomatic (non-fibrotic) convalescent and COVID-19 fibrosing interstitial lung disease patients compared to healthy controls 6 months after infection. Table showing LTA™ quantitative analysis results of CT at time of diagnosis, stratified by COVID-19 convalescent disease status. 6 healthy controls, 13/17 symptomatic and 17/17 COVID-19 FILD patients had suitable CT scans available. Statistical significance (p<0.05) was determined by one way ANOVA.

**Table S3:** Cohort 2 (convalescent non-fibrotic COVID-19) and Cohort 3 (pulmonary fibrosis) patient characteristics. Table showing demographics, previous medical history, follow up symptoms and inpatient severity of convalescent patients hospitalised with COVID-19 (cohort 2) and patients with COVID-19 fibrosing interstitial lung disease (fILD) and idiopathic pulmonary fibrosis (IPF). Abbreviations and units: M, male; F, female; BMI, body mass index, kg/m^2^; CRP, C-reactive protein, mg/L; LOS, length of stay, days. Statistical significance (p<0.05) was determined using one way ANOVA or Χ^2^ testing as appropriate. *p values represent statistical significance between asymptomatic and symptomatic COVID-19 convalescent groups. **p values represent statistical significance between COVID-19 FILD and IPF groups. ^Ϯ^pulmonary diseases excluding asthma, which is presented separately. Values represent mean and () standard deviation of the mean.

|  |  | Cohort 2 |  |  | Cohort 3 |  |  |
| --- | --- | --- | --- | --- | --- | --- | --- |
|  |  | Asymptomatic<br>convalescent<br>(n=9) | Symptomatic<br>convalescent<br>(n=17) | p<br>value*<br>(**<0.05**) | COVID-19<br>ILD<br>(n=17) | IPF<br>(n=12) | P<br>value**<br>(**<0.05**) |
| Demographics | Age | 60.0<br>(15.2) | 60.3<br>(10.4) | NS | 61.4<br>(11.2) | 75.2<br>(5.8) | <0.001 |
|  | Sex<br>(M:F) | 3.5 | 4.25 | NS | 16.0 | 11.0 | NS |
|  | Ethnicity<br>(Caucasian%) | 70.0 | 76.5 | NS | 64.7 | 100.0 | 0.02 |
|  | BMI | 29.4<br>(4.8) | 30.9<br>(6.2) | NS | 32.2<br>(4.5) | 31.0<br>(7.3) | NS |
| Previous<br>medical<br>history | Hypertension | 66.7 | 41.2 | NS | 76.5 | 50 | NS |
|  | IHD | 22.2 | 11.8 | NS | 11.8 | 16.7 | NS |
|  | Pulmonary<sup>†</sup> | 22.2 | 0.0 | <0.05 | 5.9 | 25.0 | NS |
|  | Asthma | 11.1 | 23.5 | NS | 23.5 | 8.3 | NS |
|  | Diabetes | 22.2 | 6.3 | NS | 58.8 | 25.0 | 0.06 |
| Follow up<br>symptom | Breathless | 0.0 | 58.8 | <0.01 | 94.4 | 100.0 | NS |
|  | Cough | 0.0 | 47.1 | <0.01 | 64.7 | 75.0 | NS |
|  | Fatigue | 0.0 | 58.8 | <0.01 | 52.9 | 8.3 | 0.02 |
|  | Myalgia | 0.0 | 35.2 | <0.01 | 5.9 | 8.3 | NS |
| Inpatient<br>Severity | Mild (n) | 3 | 4 | NS | 1 |  |  |
|  | Moderate (n) | 3 | 6 |  | 1 |  |  |
|  | Severe (n) | 3 | 7 |  | 15 |  |  |
|  | CRP | 114.6<br>(76.6) | 168.4<br>(109.0) | NS | 147.1<br>(58.1) |  |  |
|  | Length of stay | 9.8<br>(7.5) | 9.8<br>(9.0) | NS | 22.3<br>(13.6) |  |  |
| Time to follow up (days) |  | 141.2<br>(43.0) | 162.0<br>(34.7) | NS | 271.1<br>(119.1) |  |  |

**Table S4:** Lung function tests do not differ between asymptomatic and (non-fibrotic) symptomatic convalescent COVID-19 patients. Table showing lung function physiology tests performed on convalescent patients hospitalised with COVID-19 (cohort 2). Statistical significance (p<0.05) tested by unpaired t-test. Units of measurement: FEV_1_, litres/min; FVC, litres; millilitres CO/minute/mm Hg. % predicted is a comparison to the GLI (2017) reference values. Values in () represent standard deviation of the mean.

|  | Lung function- stratified by resolution of symptoms of convalescent non-fibrotic COVID-19 patients |  |  |  |  |  |  |
| --- | --- | --- | --- | --- | --- | --- | --- |
|  | FEV<sub>1</sub> (L) | FEV<sub>1</sub> (%) predicted | FVC (L) | FVC (%) predicted | DLCO | DLCO (%) predicted | KCO (%) predicted |
| **Asymptomatic convalescent (n=7)** | 2.83<br>(0.95) | 89.6<br>(20.7) | 3.68<br>(1.39) | 90.3<br>(25.6) | 7.15<br>(1.36) | 87.4<br>(9.5) | 94.8<br>(8.4) |
| **Symptomatic convalescent (n=16)** | 2.80<br>(0.70) | 85.3<br>(24.9) | 3.66<br>(0.94) | 93.0<br>(15.7) | 7.25<br>(1.98) | 88.9<br>(13.1) | 106.0<br>(10.5) |
| **P value (<0.05)** | NS | NS | NS | NS | NS | NS | NS |

**Table S5:** Primary antibodies used for immunohistochemistry

| **Primary antibody target** | **Species reactivity** | **Animal raised in** | **Company (product number)** | **Concentration** |
| --- | --- | --- | --- | --- |
| **SOX9** | human/mouse | Rabbit | Abcam 185966 | 1:1000 |
| **COL1** | human/mouse | Goat | Southern bio 1310-01 | 1:1000 |
| **α-SMA** | human/mouse | Mouse | Leica NCL-L-SMA | 1:100 |
| **PDGFR-β** | human/mouse | Rabbit | Abcam 32570 | 1:400 |
| **CD31** | human/mouse | Rabbit | Abcam 28364 | 1:100 |
| **Pro-SFTPC** | human/mouse | Rabbit | Abcam 90716 | 1:1000 |
| **SPRR1a** | human/mouse | Rabbit | Thermofisher PA5-26568 | 1:250 |
| **KRT17** | human/mouse | Rabbit | Abcam 53707 | 1:500 |

**Table S6:** Secondary antibodies used for immunohistochemistry

| **Secondary antibody target** | **Species reactivity** | **Animal raised in** | **Company (product number)** | **Concentration** |
| --- | --- | --- | --- | --- |
| **Rabbit Biotinylated** | Rabbit | Goat | Vector BA-1000 | 1:800 |
| **Goat Biotinylated** | Goat | Horse | Vector BA-9500 | 1:500 |
| **Mouse Biotinylated** | Mouse | Goat | Vector BA-9200 | 1:800 |
| **ImPRESS<sup>TM</sup> anti- rabbit HRP** | Rabbit | Horse | Vector MP-7401 | Ready to use |
| **ImPRESS<sup>TM</sup> anti- mouse HRP** | Mouse | Horse | Vector MP-7402 | Ready to use |
| **ImPRESS<sup>TM</sup> anti- rabbit AP** | Rabbit | Horse | Vector MP-5401 | Ready to use |
| **ImPRESS<sup>TM</sup> anti- mouse AP** | Mouse | Horse | Vector MP-5402 | Ready to use |

**Table S7:**
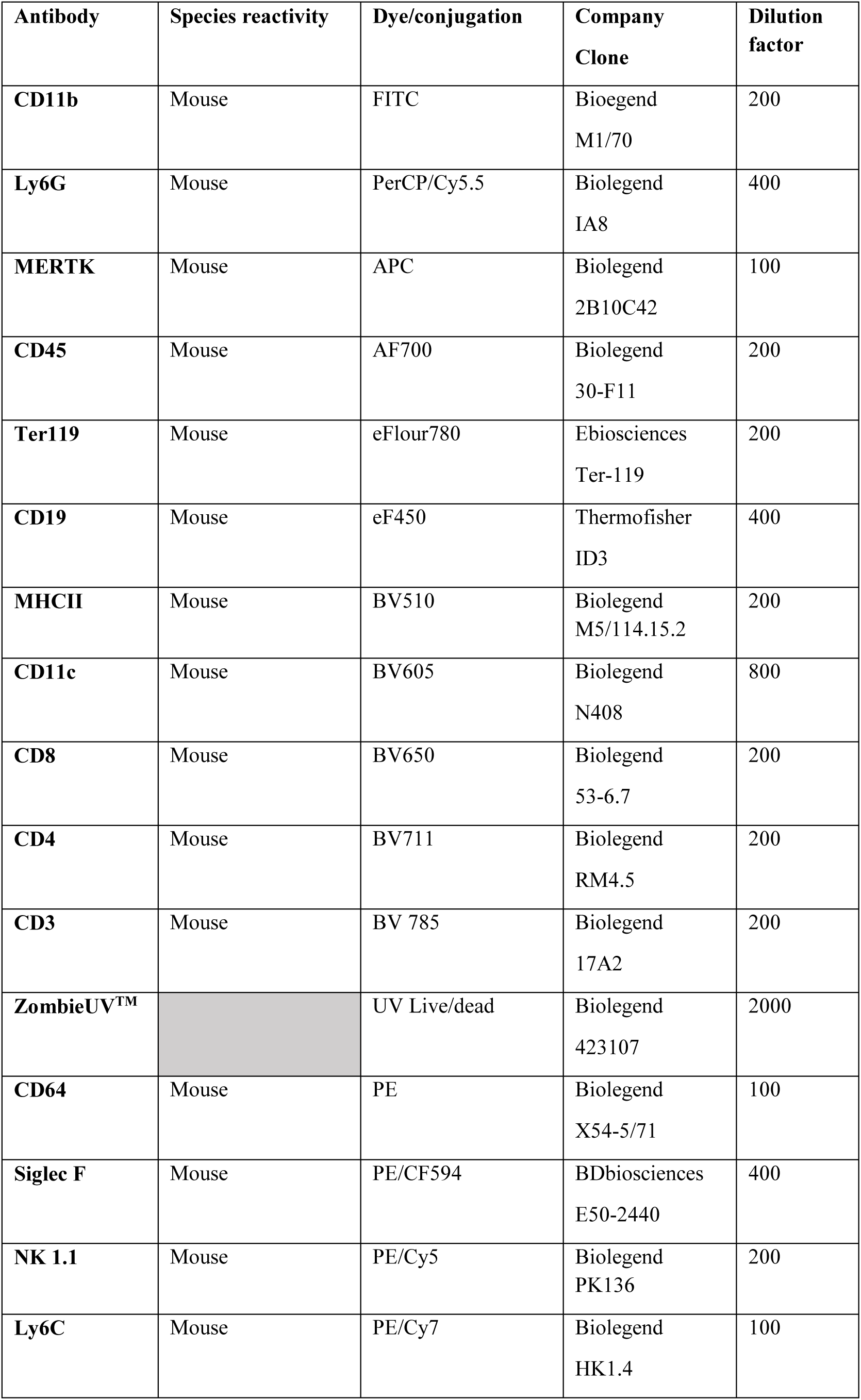
Flow cytometry antibodies

| **Antibody** | **Species reactivity** | **Dye/conjugation** | **Company**<br>**Clone** | **Dilution**<br>**factor** |
| --- | --- | --- | --- | --- |
| **CD11b** | Mouse | FITC | Bioegend<br>M1/70 | 200 |
| **Ly6G** | Mouse | PerCP/Cy5.5 | Biolegend<br>IA8 | 400 |
| **MERTK** | Mouse | APC | Biolegend<br>2B10C42 | 100 |
| **CD45** | Mouse | AF700 | Biolegend<br>30-F11 | 200 |
| **Ter119** | Mouse | eFlour780 | Ebiosciences<br>Ter-119 | 200 |
| **CD19** | Mouse | eF450 | Thermofisher<br>ID3 | 400 |
| **MHCII** | Mouse | BV510 | Biolegend<br>M5/114.15.2 | 200 |
| **CD11c** | Mouse | BV605 | Biolegend<br>N408 | 800 |
| **CD8** | Mouse | BV650 | Biolegend<br>53-6.7 | 200 |
| **CD4** | Mouse | BV711 | Biolegend<br>RM4.5 | 200 |
| **CD3** | Mouse | BV 785 | Biolegend<br>17A2 | 200 |
| **ZombieUV™** |  | UV Live/dead | Biolegend<br>423107 | 2000 |
| **CD64** | Mouse | PE | Biolegend<br>X54-5/71 | 100 |
| **Siglec F** | Mouse | PE/CF594 | BDbiosciences<br>E50-2440 | 400 |
| **NK 1.1** | Mouse | PE/Cy5 | Biolegend<br>PK136 | 200 |
| **Ly6C** | Mouse | PE/Cy7 | Biolegend<br>HK1.4 | 100 |

**Table S8:** Western blot antibodies

| **Primary antibodies** |  |  |  |  |
| --- | --- | --- | --- | --- |
| **Antibody target** | Reactivity | Animal raised in | Company (product no.) | Dilution |
| **SOX9** | Human/mouse | Rabbit | Abcam<br>185966 | 1:1000 |
| **COL1** | Human/mouse | Goat | Southern bio<br>1310-01 | 1:3000 |
| **α-SMA** | Human/mouse | Mouse | Leica<br>NCL-L-SMA | 1:250 |
| **Secondary antibodies** |  |  |  |  |
| **Rabbit** | Rabbit | Donkey | GE Healthcare<br>(NA934) | 1:10,000 |
| **Goat** | Goat | Rabbit | GE healthcare<br>(P0449) | 1:2000 |
| **Mouse** | Mouse | Sheep | GE Healthcare<br>(NXA931) | 1:25,000 |

**Table S9:**
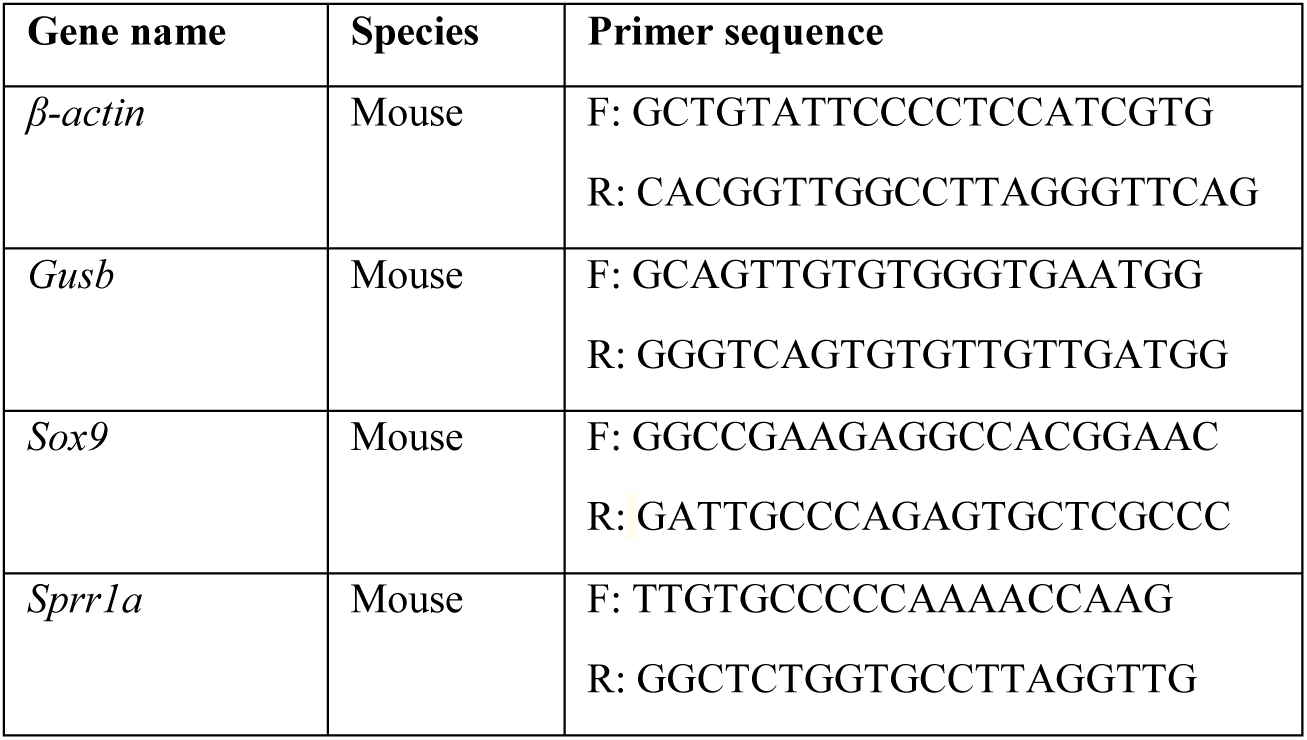
Quantitative polymerase chain reaction (qPCR) primer sequences.

